# The SCP4-STK35/PDIK1L complex is a dual phospho-catalytic signaling dependency in acute myeloid leukemia

**DOI:** 10.1101/2021.05.09.443327

**Authors:** Sofya A. Polyanskaya, Rosamaria Y. Moreno, Bin Lu, Ruopeng Feng, Yu Yao, Seema Irani, Olaf Klingbeil, Zhaolin Yang, Yiliang Wei, Osama E. Demerdash, Lukas A. Benjamin, Mitchell J. Weiss, Yan Jessie Zhang, Christopher R. Vakoc

## Abstract

Acute myeloid leukemia (AML) cells rely on phospho-signaling pathways to gain unlimited proliferation potential. Here, we used domain-focused CRISPR screening to identify the nuclear phosphatase SCP4 as a dependency in AML, yet this enzyme is dispensable in normal hematopoietic progenitor cells. Using CRISPR exon scanning and gene complementation assays, we show that the catalytic function of SCP4 is essential in AML. Through mass spectrometry analysis of affinity-purified complexes, we identify the kinase paralogs STK35 and PDIK1L as binding partners and substrates of the SCP4 phosphatase domain. We show that STK35 and PDIK1L function catalytically and redundantly in the same pathway as SCP4 to maintain AML proliferation and to support amino acid biosynthesis and transport. We provide evidence that SCP4 regulates STK35/PDIK1L through two distinct mechanisms: catalytic removal of inhibitory phosphorylation and by promoting kinase stability. Our findings reveal a phosphatase-kinase signaling complex that supports the pathogenesis of AML.

## INTRODUCTION

Acute myeloid leukemia (AML) is an aggressive hematopoietic malignancy caused by the impaired differentiation and unrestrained proliferation of myeloid progenitor cells (Grove and Vassiliou, 2014). While genetically heterogeneous, most cases of AML harbor a mutation that deregulates signaling downstream of growth factor receptors (Bullinger et al., 2017). For example, activating mutations in the receptor tyrosine kinase FLT3 are present in one-third of AML cases, and inhibitors of this kinase are an approved therapy in this group of patients (Daver et al., 2019; Short et al., 2019; Kim, 2017; Dhillon, 2019). An example of a phosphatase target in leukemia is the oncoprotein SHP2, which is a positive regulator of RAS/MAP kinase signaling and can be targeted by allosteric small-molecule inhibitors (Elson, 2018; Yuan et al., 2020; Vainonen et al., 2021). Based on the actionability of kinase and phosphatase targets, a major goal in AML research is to discover and validate acquired dependencies on phospho-catalytic signaling proteins using functional genomics.

SCP4 (encoded by *CTDSPL2*) is one of the least characterized human phosphatases, with yet-to-be-determined roles in human cancer. SCP4 belongs to the haloacid dehalogenase (HAD) family of proteins, which employ conserved active site aspartates within a DxDx(V/T) motif to remove phosphates from protein or non-protein substrates (Seifried et al., 2013). SCP4 localizes to the nucleus, where it can dephosphorylate several putative substrates, including transcription factors and RNA polymerase II (Zhao et al., 2014; Wani et al., 2016; Liu et al., 2017; Zhao et al., 2018). SCP4^-/-^ mice have been characterized and found to complete embryonic development, but these animals die shortly after birth due to defective gluconeogenesis (Cao et al., 2018). This phenotype was linked to the deregulation of FoxO transcription factors, suggesting that a critical function of SCP4 is to regulate metabolism (Cao et al., 2018). In chicken models of cancer, the *CTDSPL2* ortholog is a common integration site for Avian leukosis virus-induced B-cell lymphomas (Winans et al., 2017). Despite belonging to a potentially druggable family of proteins (Krueger et al., 2014), it has yet to be determined whether any human cancers rely on SCP4 for disease initiation or progression.

Another set of poorly characterized signaling proteins are the kinase paralogs STK35 and PDIK1L, which share 69% sequence identity. STK35 and PDIK1L localize to the nucleus, but the function of these enzymes in this cellular compartment is unclear. A small number of studies suggest that STK35 can regulate cell proliferation, apoptosis, and metabolism, albeit via unclear molecular mechanisms (Wu et al., 2018; Yang et al., 2020). In addition, the expression of STK35/PDIK1L correlates with disease progression and prognosis in several tissue contexts (Wu et al., 2018; Yang et al., 2020; Greenbaum et al., 2021). Nonetheless, the signaling function and biochemical interactions of STK35/PDIK1L remain unknown.

In this study, we used unbiased genetic and biochemical approaches to show that SCP4 forms a dual catalytic signaling complex with STK35/PDIK1L to support the pathogenesis of AML. Within this complex, SCP4 is critical for maintaining STK35/PDIK1L stability and also acts to remove inhibitory phosphorylation from the kinase activation loop. A downstream output of this phospho-catalytic complex in AML is to maintain the expression of genes involved in amino acid biosynthesis and transport. Taken together, our study identifies a nuclear SCP4-STK35/PDIK1L signaling complex and reveals its critical role in the pathogenesis of AML.

## RESULTS

### Phosphatase domain-focused CRISPR screening identifies context-specific dependencies in human cancer cell lines

We previously described a domain-focused CRISPR-Cas9 genetic screening strategy that can nominate catalytic dependencies in cancer by targeting indel mutagenesis to domain-encoding exons (Shi et al., 2015). Here, we applied this method to the human phosphatase protein family in search of AML-specific dependencies. For this purpose, we designed and cloned a pooled library of 1,961 sgRNAs targeting exons that encode 217 known or putative phosphatase domains, including phosphatases with protein and non-protein substrates (Figure 1A).

**Figure 1.**
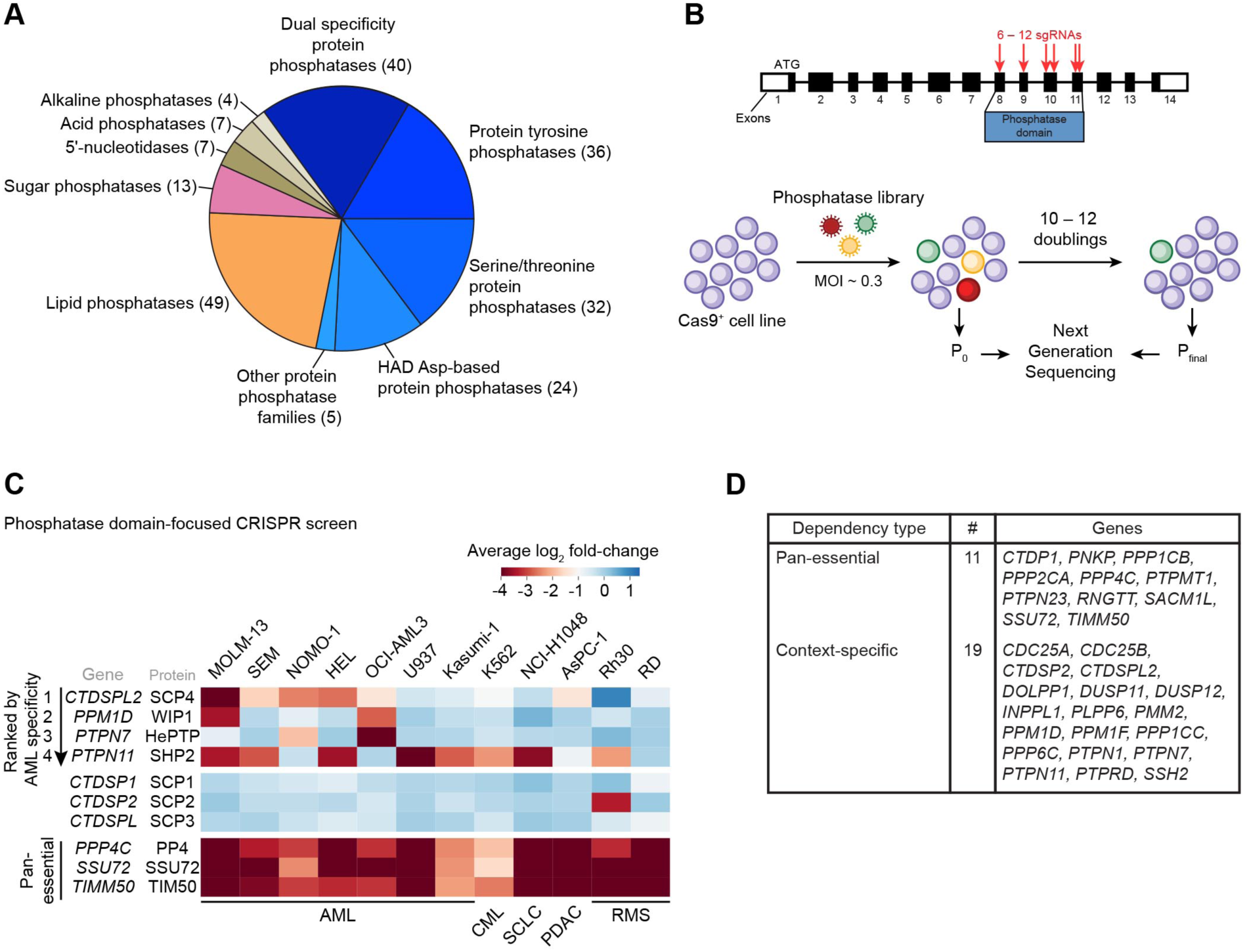
Phosphatase domain-focused CRISPR screening identifies context-specific dependencies in human cancer cell lines. **(A)** Categorization of human phosphatase domains targeted by CRISPR-Cas9 genetic screens in this study. **(B)** Overview of the genetic screening procedure using the phosphatase domain-focused sgRNA library. **(C)** Extracted essentiality scores for phosphatases demonstrating AML-bias or pan-essentiality. Plotted is the log_2_ fold-change of sgRNA abundance during ∼11 population doublings. The effects of individual sgRNAs targeting each domain were averaged. AML, acute myeloid leukemia; CML, chronic myeloid leukemia; SCLC, small cell lung cancer; PDAC, pancreatic ductal adenocarcinoma; RMS, rhabdomyosarcoma. **(D)** Summary of dependencies discovered in the screen. See also Figure S1.

We used this library to perform negative-selection ‘dropout’ screens in eight leukemia and four solid tumor cell lines (Figure 1B, 1C, S1A). The performance of spike-in negative and positive control sgRNAs in this library validated the overall quality of these screens (Figure S1B). This screen revealed that sgRNAs targeting 187 of the phosphatase domains showed no significant impairment of cell fitness in any of the cell lines tested, whereas 11 phosphatases were pan-essential across the entire panel of cell lines (Figure 1D and S1A, Table S1). The remainder showed varying levels of specificity, which included the validation of known context-specific phosphatase dependencies: SHP2 was only essential in cell lines that lacked activating mutations of RAS (Ahmed et al., 2019), PPM1D was only required in TP53 wild-type lines (Goloudina et al., 2016), and PTP1B was only a dependency in a BCR-ABL1 positive leukemia cell line (Alvira et al., 2011) (Figure S1C-S1E).

Next, we ranked all of the phosphatases based on their relative essentiality in AML versus non-AML contexts, which nominated SCP4 as the top-ranking AML-biased dependency (Figure 1C and S1A). Among the AML cell lines screened, five showed SCP4 dependency, with MOLM-13 being the most sensitive. In contrast, AML cell lines U937 and Kasumi-1 and the CML line K562 did not show significant cell fitness defects upon SCP4 targeting. The closest paralogs of SCP4 were not essential in AML (Figure 1C). Since no prior study has implicated SCP4 in human cancer, we investigated further the essential function of this phosphatase in AML.

### SCP4 is an acquired dependency in human AML cells

To validate the AML-biased essentiality of SCP4, we performed competition-based proliferation assays comparing SCP4 sgRNAs to positive and negative control sgRNAs in a total of 23 human cancer cell lines: 14 leukemia, 4 pancreatic cancer, 3 rhabdomyosarcoma, and 2 lung cancer lines (Figure 2A). In these experiments, ten leukemia cell lines demonstrated a clear dependency on SCP4, while four of the leukemia lines and all of the solid tumor cell lines were insensitive to SCP4 sgRNAs. The sensitive cell lines included engineered human MLL-AF9/Nras^G12D^ and MLL-AF9/FLT3^ITD^ AML cell lines (Wei et al., 2008). In contrast, an sgRNA targeting the DNA replication protein PCNA suppressed the growth of all cell lines tested. Western blot analysis confirmed that the differential SCP4 requirement was not because of differences in genome editing efficiency (Figure 2B-2E). Rescue experiments using an sgRNA-resistant cDNA confirmed that the growth arrest caused by SCP4 sgRNAs occurred via an on-target mechanism (Figure S2A-S2C). Using flow cytometry-based BrdU incorporation measurements and Annexin V staining, we found that SCP4 inactivation in MOLM-13 cells led to a G1/G0-arrest and the induction of apoptosis (Figure 2F, S2D, and S2E). We also validated that targeting SCP4 attenuated the growth of MOLM-13 cells when injected via tail vein and expanded in immune-deficient mice (Figure 2G–2H). As additional validation, we verified that Scp4 was essential in an engineered murine AML cell line MLL-AF9/Nras^G12D^ (RN2) (Zuber et al., 2011), an effect that was rescued by the expression of the human SCP4 cDNA (Figure S2F). However, immortalized NIH3T3 murine fibroblasts were unaffected by SCP4 sgRNAs (Figure S2G), suggesting that the AML-biased dependency on SCP4 is conserved across species.

**Figure 2.**
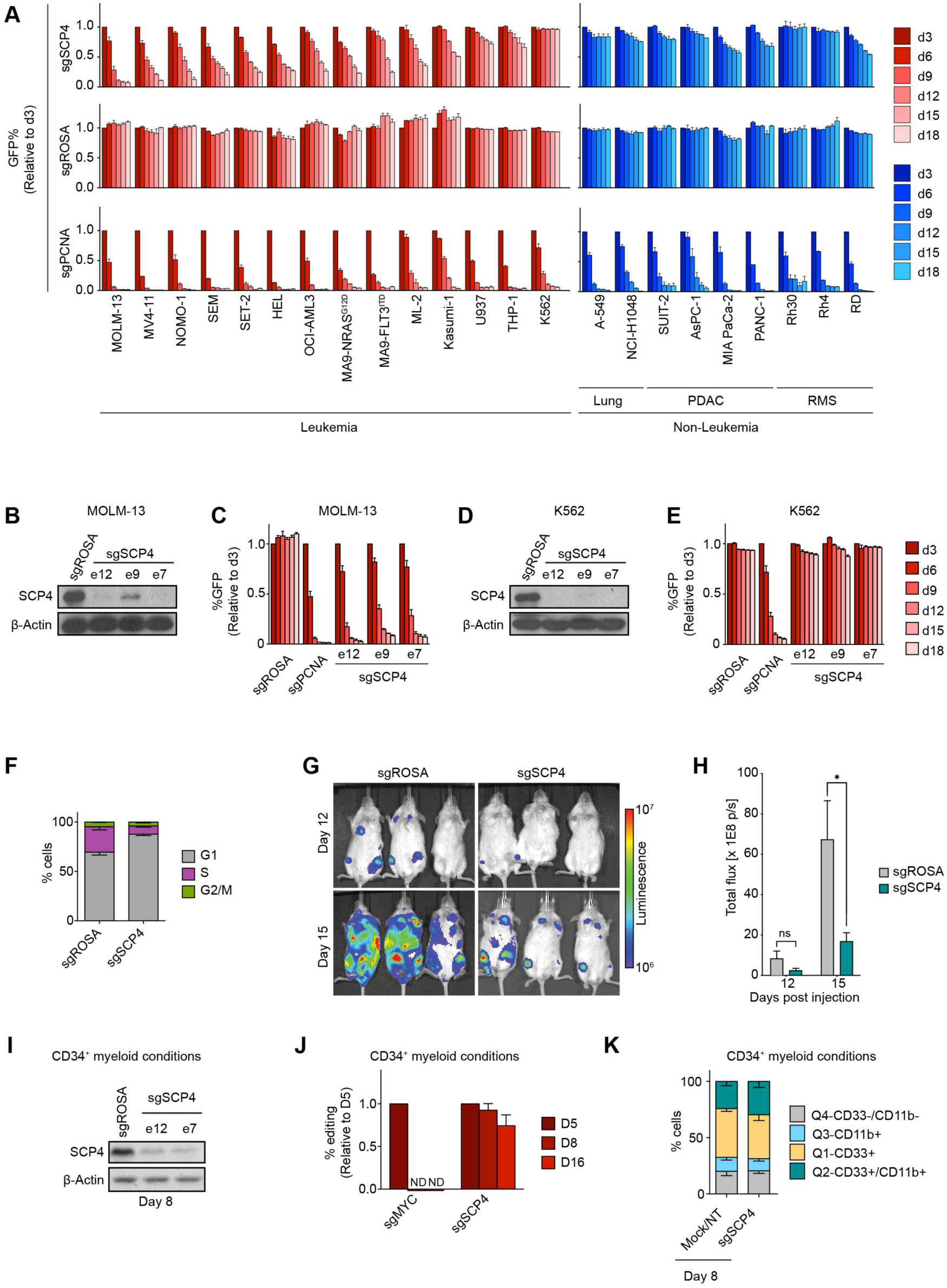
SCP4 is an acquired dependency in human AML. **(A)** Competition-based proliferation assays in 23 human cancer cell lines infected with the indicated sgRNAs. PDAC, Pancreatic Ductal Adenocarcinoma; RMS, rhabdomyosarcoma. n=3. **(B)** Western blot of whole-cell lysates from MOLM-13 cells on day 5 post-infection with the indicated sgRNAs. **(C)** Competition-based proliferation assay in MOLM-13 cells infected with the indicated sgRNAs. n=3. **(D)** Western blot of whole-cell lysates from K562 cells on day 5 post-infection with the indicated sgRNAs. **(E)** Competition-based proliferation assay in K562 cells infected with the indicated sgRNAs. n=3. **(F)** Quantification of the different cell cycle stages in MOLM-13 cells on day 5 post-infection with the indicated sgRNAs. n = 3. **(G)** Bioluminescence imaging of NSG mice transplanted with luciferase^+^/Cas9^+^ MOLM-13 cells infected with either sgROSA or sgSCP4. **(H)** Quantification of bioluminescence intensity. n=3. p-value was calculated by unpaired Student’s t-test. *p < 0.05. **(I)** Western blot of whole-cell lysates from CD34^+^ cells electroporated with Cas9 loaded with the indicated sgRNAs, day 8 post-electroporation, culturing in myeloid conditions. **(J)** RT-qPCR analysis of indels presence in CD34^+^ cells electroporated with Cas9 loaded with the indicated sgRNAs over the course of culturing in myeloid conditions. The effects of individual sgRNAs for SCP4 were averaged. n = 4. **(K)** Quantification of the flow cytometry analysis of myeloid differentiation of CD34^+^ cells electroporated with Cas9 loaded with the indicated sgRNAs, day 8 post-electroporation, culturing in myeloid conditions. The effects of individual negative controls and sgRNAs for SCP4 were averaged. n = 4. All bar graphs represent the mean ± SEM. All sgRNA experiments were performed in Cas9-expressing cell lines. ‘e’ refers to the exon number targeted by each sgRNA. The indicated sgRNAs were linked to GFP. GFP^+^ population depletion indicates loss of cell fitness caused by Cas9/sgRNA-mediated genetic mutations. ROSA, Mock, and NT, negative controls; PCNA and MYC, positive controls. See also Figure S2.

To evaluate the SCP4 requirement in non-transformed hematopoietic cells, we performed electroporation of Cas9:gRNA ribonucleoprotein complexes into human CD34^+^ hematopoietic stem and progenitor cells (HSPCs) (Metais et al., 2019). After electroporation, successful inactivation of SCP4 was verified by western blotting and by measuring the presence of *CTDSPL2* indel mutations (Figure 2I-2J, S2H-S2L). Unlike the effects of targeting the essential transcription factor MYC, we observed no evidence that SCP4-deficient cells experienced any fitness disadvantage during 16 days of culturing under conditions that promote myeloid, erythroid, or megakaryocytic differentiation (Figure 2J, S2K, and S2L). In addition, flow cytometry analysis and methylcellulose plating experiments showed that SCP4-deficient HSPCs showed no defect in differentiation into these three lineages (Figure 2K and S2M-S2O). These findings are consistent with the survival of SCP4^-/-^ mice to the neonatal stage of development (Cao et al., 2018) since a severe defect in hematopoiesis would have led to the embryonic lethality of these animals. Collectively, these data suggest that SCP4 is an acquired dependency in AML.

### The catalytic phosphatase activity of SCP4 is essential in AML

The C-terminal phosphatase domain of SCP4 (amino acids 236–442) is highly conserved among vertebrates, whereas amino acids 1–236 are less conserved and are predicted to be intrinsically disordered (Figure 3A and 3B) (Chang et al., 2018; Meszaros et al., 2018). In an initial effort to map the functionally important regions of SCP4 in AML, we performed CRISPR exon scanning (Shi et al., 2015). We constructed a pooled library of 85 sgRNAs tiling along each of the *CTDSPL2* exons and used this for negative selection CRISPR-Cas9 screening in MOLM-13 cells. By quantifying sgRNA depletion along the length of the gene, we observed a pattern in which sgRNAs targeting exons encoding amino acids 1–235 resulted in a minimal growth arrest phenotype, whereas sgRNAs targeting the 236–466 segment resulted in significant negative selection (Figure 3C, Table S2). This result suggests that indel mutations produce a higher proportion of null alleles in the C-terminal segment of SCP4, which implicates the phosphatase domain as critical for AML growth (Shi et al., 2015). Notably, SCP4/*CTDSPL2* sgRNAs used in Project Achilles/DepMap genome-wide screens (Meyers et al., 2017; Dempster et al., 2019) exclusively target the 1–235 segment of the protein (Figure S3A), which explains why an AML dependency was not observed in these data.

**Figure 3.**
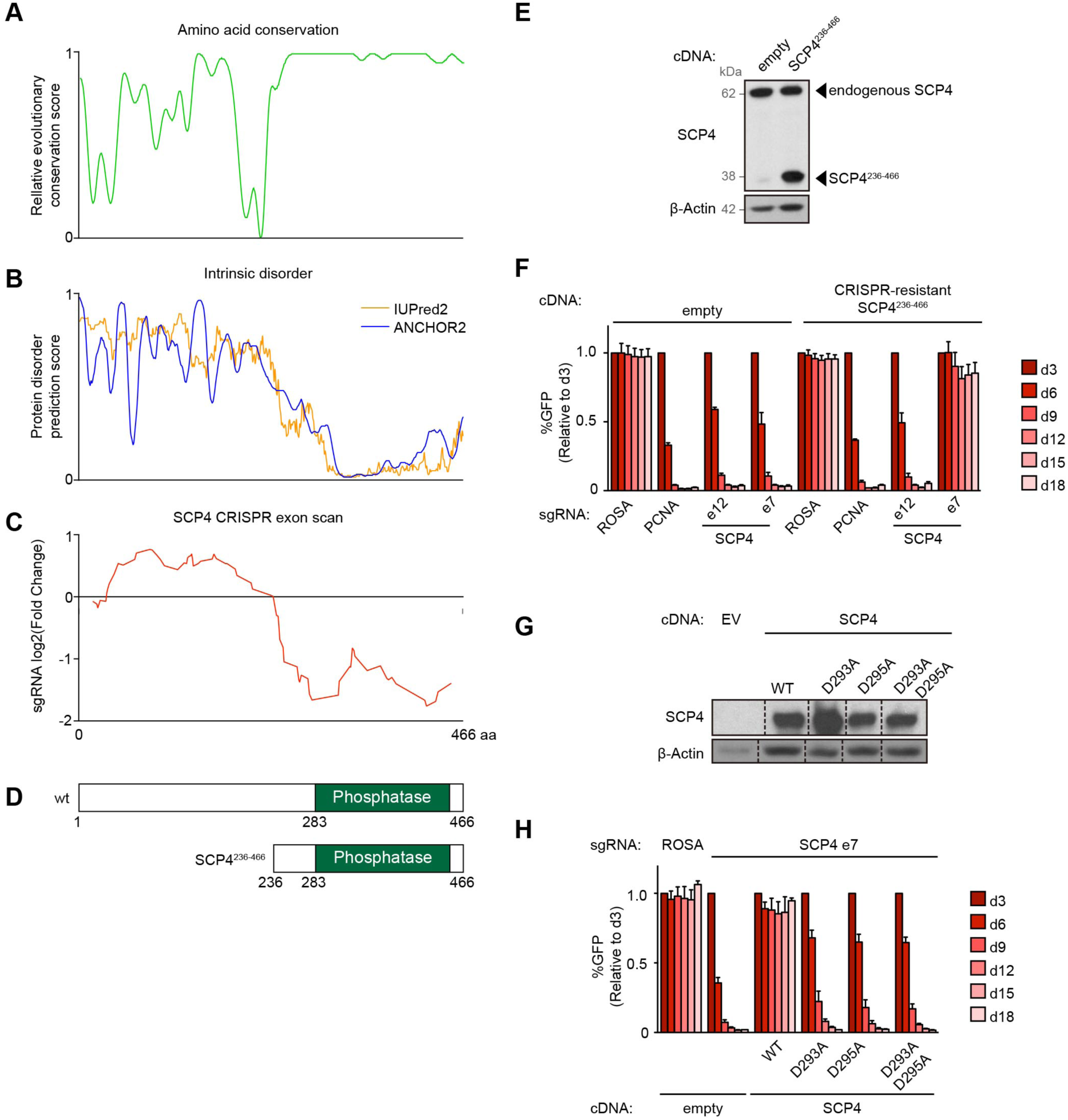
The catalytic phosphatase function of SCP4 is essential in AML. **(A)** Relative evolutionary conservation score for each residue, from 0 — least conserved to 1 — most conserved. Based on data from Chang et al., 2018. **(B)** Protein disorder prediction score for each residue, from 0 — least disordered to 1 — most disordered. Based on data from Meszaros et al., 2018. **(C)** Running average of log_2_ fold changes of the CRISPR-scan of SCP4 with all the possible sgRNAs. SCP4 protein amino acid numbers are indicated along the x axis. **(D)** Domain architectures of human SCP4 and the truncated version of SCP4 used in this study. **(E)** Western blot of whole-cell lysates from MOLM-13 cells stably expressing empty vector or CRISPR-resistant SCP4^236-466^. **(F)** Competition-based proliferation assay in MOLM-13 cells stably expressing empty vector or CRISPR-resistant SCP4^236-466^ infected with the indicated sgRNAs. n = 3. **(G)** Western blot of whole-cell lysates from MOLM-13 cells stably expressing empty vector (EV, underloaded) or CRISPR-resistant wild type (WT) and catalytic mutants of SCP4. Vertical black dashed lines indicate omitted lanes from the same gel, same exposure. **(H)** Competition-based proliferation assay in MOLM-13 cells stably expressing empty vector or CRISPR-resistant wild type (WT) and catalytic mutants of SCP4 infected with the indicated sgRNAs. n = 3. All bar graphs represent the mean ± SEM. All sgRNA experiments were performed in Cas9-expressing cell lines. ‘e’ refers to the exon number targeted by each sgRNA. The indicated sgRNAs were linked to GFP. GFP^+^ population depletion indicates loss of cell fitness caused by Cas9/sgRNA-mediated genetic mutations. ROSA, negative control; PCNA, positive control. See also Figure S3.

To corroborate the CRISPR exon scanning results, we performed cDNA rescue experiments evaluating the ability of different SCP4 truncations and mutations to complement the inactivation of endogenous SCP4. Similar to results obtained with the wild-type protein, the SCP4^236-466^ truncation was sufficient to rescue the knockout of endogenous *CTDSPL2*, verifying that the N-terminal half of SCP4 is dispensable in AML (Figure 3D-3F). In addition, we cloned point mutations of SCP4 that changed aspartate residues in the putative DxDx(V/T) catalytic motif to alanine. Despite similar expression levels to the wild-type protein, SCP4^D293A^, SCP4^D295A^, and SCP4^D293A/D295A^ mutants were unable to support MOLM-13 growth (Figure 3G and 3H). Collectively, these genetic experiments suggest that the catalytic phosphatase function of SCP4 is essential in AML.

### STK35 and PDIK1L bind selectively to the catalytically active form of SCP4

In accord with prior studies (Kemmer et al., 2006; Zhao et al., 2014; Kang et al., 2016), we found that SCP4 was localized to the nucleus of AML cells (Figure 4A). However, our efforts at performing SCP4 ChIP-seq in MOLM-13 cells failed to identify evidence of sequence-specific DNA occupancy (data not shown). Therefore, we turned to biochemical approaches to further understand the mechanism of SCP4 dependency. We stably expressed a FLAG-tagged SCP4 via lentivirus in MOLM-13 cells at levels comparable to endogenous SCP4 (Figure 4B). Importantly, this epitope-tagged protein localized to the nucleus and was able to rescue the knockout of endogenous *CTDSPL2* (Figure S4A and S2B-S2C). Thereafter, we optimized an anti-FLAG affinity purification procedure that efficiently recovered FLAG-SCP4 complexes and removed endogenous SCP4 (Figure 4B). In these experiments, we compared wild-type SCP4 to SCP4^D293A^, with the intent that any protein-protein interaction abrogated by this loss-of-function mutation would have a higher likelihood of functional significance.

**Figure 4.**
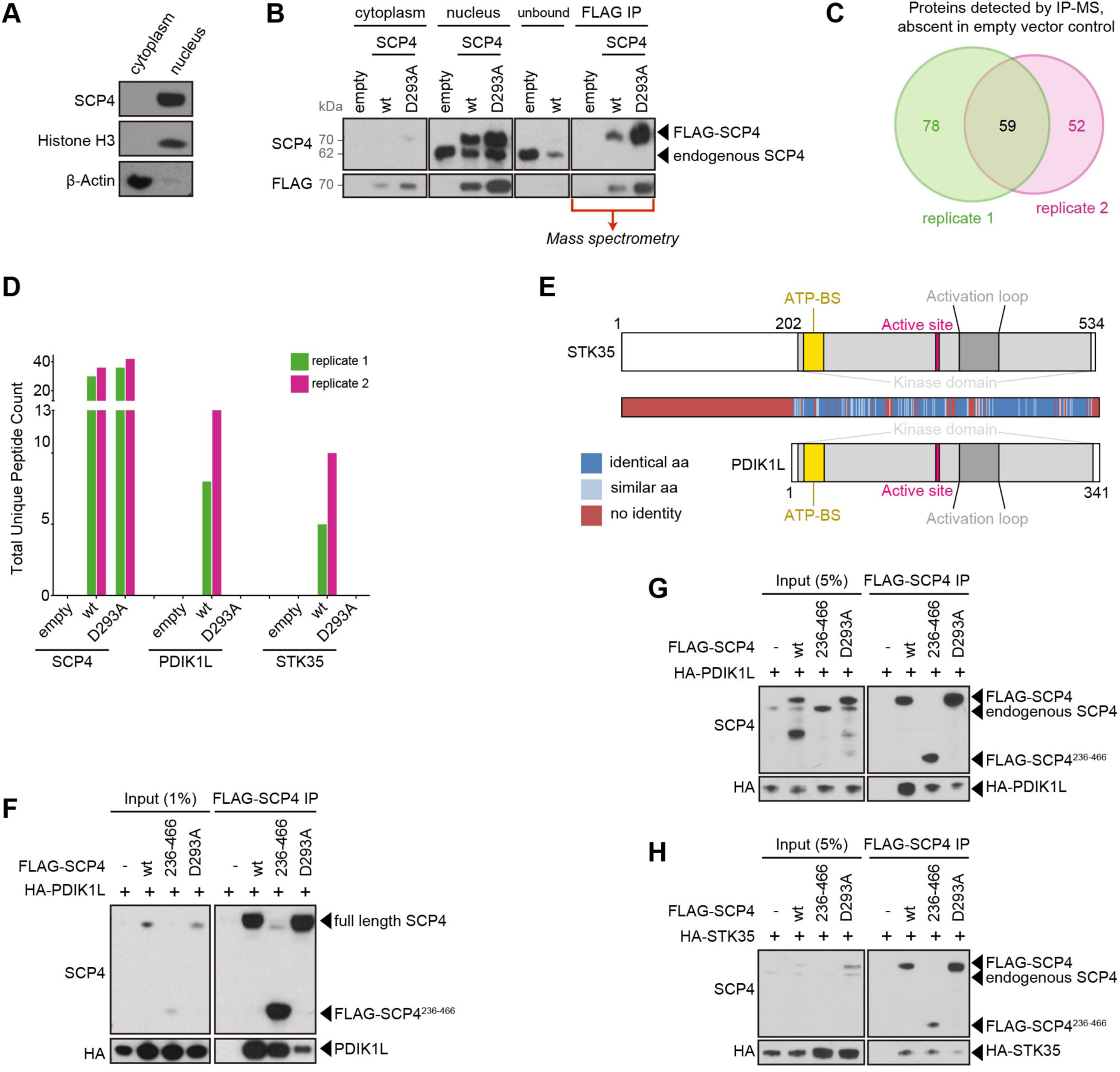
STK35 and PDIK1L bind selectively to the catalytically active form of SCP4. **(A)** Western blot of cytoplasm and nuclear fractions from MOLM-13 cells. **(B)** Representative FLAG-SCP4 affinity purification Western blot analysis for the subsequent mass spectrometry (MS) analysis. Cytoplasm and nucleus fractions from MOLM-13 cells stably expressing empty vector, FLAG-SCP4 wild type (WT), or catalytic mutant (D293A). Nuclear fraction was mixed with anti-FLAG M2 agarose overnight. The flow-through was analyzed to ensure efficient binding of the FLAG-tagged constructs (loaded as “unbound”). Following the extensive washing, the agarose amount equivalent to the cytoplasm and nucleus fractions loading was boiled in Laemmli buffer and loaded as “FLAG-IP”. The rest was sent for the MS analysis. **(C)** Venn diagram depicting the overlap between proteins detected by MS and absent in empty vector controls in the two independent biological replicates. **(D)** Total unique peptide counts for SCP4, PDIK1L, and STK35, detected by MS in the two independent biological replicates. **(E)** Domain architectures and homology heat-map of human STK35 and PDIK1L. ATP-BS, ATP-binding site. **(F–H)** Immunoprecipitation followed by Western blotting performed with the indicated antibodies. The whole-cell lysate was prepared from HEK293T 24 hours post-transfection with the indicated constructs (F). The nuclear lysates were prepared from the human MOLM-13 cells stably expressing the indicated constructs (G, H). ‘-’, empty vector; WT, wild type FLAG-SCP4; 236-466, FLAG-SCP4^236-466^; D293A, catalytic mutant FLAG-SCP4^D293A^, IP, immunoprecipitation. Note: degradation bands appear in the WT and D293A input at ∼50 kDa and in D293A at ∼40 kDa. See also Figure S4.

Mass spectrometry (MS) analysis of purified FLAG-SCP4 was performed to identify the constituents of its protein complex. A comparison of two independent biological replicates revealed 59 protein candidates (Figure 4C). Among the most abundant SCP4-associated proteins, based on total spectral counts, were several members of the importin protein family, which function to shuttle protein cargo into the nucleus (Cagatay and Chook, 2018) (Figure S4B). Our mapping experiments showed that the importin interaction occurred via the N-terminal segment 1–236, which contains predicted importin-binding nuclear localization sequences (Figure S4C-S4E). This result suggests that a function of the SCP4 N-terminal segment is to promote nuclear localization of the full-length protein. Interestingly, the low molecular weight of the SCP4^236-466^ fragment (∼38 kDa) would be expected to bypass the importin requirement for passage through the nuclear pore (Weis, 2003). This would explain the presence of SCP4^236-466^ throughout the nucleus and cytoplasm and may account for the ability of this protein to complement the knockout of endogenous *CTDSPL2* (Figure S4A and S4E).

Among the SCP4-associated proteins identified in this analysis, the paralog kinases STK35 and PDIK1L were remarkable as they were only detected in association with wild-type SCP4 but not with SCP4^D293A^ (Figure 4D and 4E). As no commercially available immune reagents were able to detect these two poorly studied proteins (data not shown), we cloned HA-tagged proteins for further biochemical analysis. Upon transient transfection in HEK293T cells or using the stable expression in MOLM-13, we found that SCP4^WT^ and SCP4^236-466^ each efficiently associated with HA-STK35 and HA-PDIK1L, whereas this interaction was diminished with the SCP4^D293A^ protein (Figure 4F-4H). This interaction was also confirmed using reciprocal IP of HA-tagged STK35/PDIK1L with endogenous SCP4 (Figure S4F-4H). Moreover, an MS analysis of immunopurified HA-PDIK1L complexes from MOLM-13 cells following high-salt washes identified endogenous SCP4 as the most enriched protein with comparable spectral counts to the PDIK1L bait (Figure S4I). These biochemical experiments suggest that the kinases STK35 and PDIK1L exist in a stable complex with the catalytically active SCP4 phosphatase domain.

### STK35 and PDIK1L function redundantly in the same genetic pathway as SCP4

We previously performed kinase domain-focused CRISPR screening in human cancer cell lines, which did not reveal a robust dependency on STK35 or PDIK1L in AML cell lines (Tarumoto et al., 2018). Since both kinases are expressed in AML (Figure 4D, S4B, and S6A), we hypothesized that functional redundancy might conceal their essentiality in this context. We investigated this by transducing human cancer cell lines with STK35 and PDIK1L sgRNAs, alone and in combination, and performing competition-based proliferation assays. These experiments revealed that MOLM-13 cells arrested their proliferation in response to the dual targeting of STK35 and PDIK1L, but the cells were less sensitive to targeting each gene individually (Figure 5A). In contrast, K562 cells did not show any phenotype following co-transduction with STK35/PDIK1L sgRNAs (Figure 5B). The proliferation-arrest phenotype of STK35/PDIK1L double knockout MOLM-13 cells was also fully rescued by expressing an sgRNA-resistant STK35 or PDI1K1L cDNA (Figure 5C). Western blotting experiments confirmed the on-target editing of the lentivirally expressed wild-type cDNA but not of the engineered sgRNA-resistant cDNA (Figure 5D). By expanding this dual targeting experiment to more leukemia lines using a bi-cistronic sgRNA vector, we found that the redundant STK35/PDIK1L dependency correlated with SCP4 dependency. For example, SCP4 and STK35/PDIK1L were not required in THP-1, U937, and K562 cells compared to the four sensitive AML cell lines (Figure 5E). These experiments demonstrate that STK35 and PDIK1L are redundant dependencies in AML and suggest that their function is linked to the SCP4 requirement in this context.

**Figure 5.**
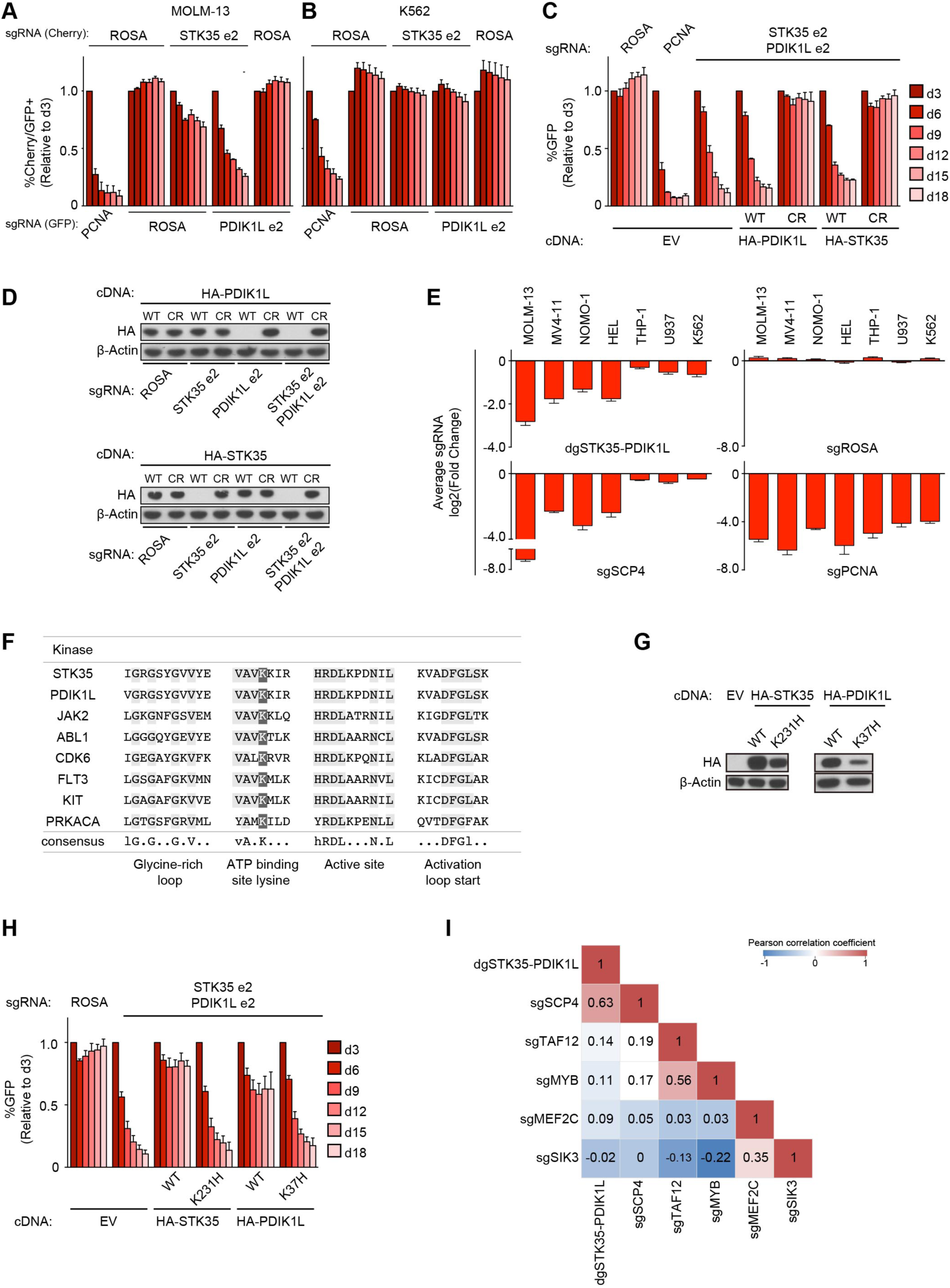
STK35 and PDIK1L function redundantly in the same genetic pathway as SCP4. **(A–B)** Competition-based proliferation assays in MOLM-13 (A) and K562 (B) cells co-infected with GFP-linked sgRNA and mCherry-linked sgRNA. Double mCherry^+^/GFP^+^ population depletion indicates loss of cell fitness due to genetic redundancy. n = 3. **(C)** Competition-based proliferation assay in MOLM-13 cells stably expressing empty vector (EV), wild type (WT), or CRISPR-resistant (CR) HA-PDIK1L and HA-STK35 infected with the indicated sgRNAs. The indicated sgRNAs were linked to GFP. n = 3. **(D)** Western blot of whole-cell lysates from MOLM-13 cells stably expressing wild type (WT) and CRISPR-resistant (CR) HA-PDIK1L and HA-STK35 on day 5 post-infection with the indicated sgRNAs. **(E)** Summary of competition-based proliferation assays in the indicated cell lines. Plotted is the fold change (log_2_) of sgRNA^+^/GFP^+^ cells after 18 days in culture (average of triplicates). **(F)** Alignment of residues defining ATP binding site for human protein kinases. Residues that are conserved among the kinases sequences are highlighted in light gray. Dark gray indicates the conserved lysine at the ATP binding site. Consensus sequence given; uppercase = conserved, lowercase = nearly invariant. The predicted structural elements of protein kinases are indicated below the consensus sequence. **(G)** Western blot of whole-cell lysates from MOLM-13 cells stably expressing empty vector (EV) or CRISPR-resistant wild type (WT) and catalytic mutants of STK35 and PDIK1L in MOLM-13. **(H)** Competition-based proliferation assay in MOLM-13 cells stably expressing empty vector (EV) or CRISPR-resistant wild type (WT) and catalytic mutant versions of HA-STK35 and HA-PDIK1L infected with the indicated sgRNAs. The indicated sgRNAs were linked to GFP. n = 3. **(I)** Correlation matrix for the global log2 fold-changes of gene expression relative to negative controls in each independent experiment across 15,095 genes. All bar graphs represent the mean ± SEM. All sgRNA experiments were performed in Cas9-expressing cell lines. ‘e’ refers to the exon number targeted by each sgRNA. Starting from Figure 5C and for the rest of the study bi-cistronic vector for simultaneous targeting of STK35 and PDIK1L was used. ‘dg’ refers to the bi-cistronic vector. ROSA, negative control; PCNA, positive control.

We next investigated whether the catalytic kinase function of STK35/PDIK1L was essential for AML cell proliferation. Sequence alignment identified the conserved ATP-binding lysine residue present in other catalytically active kinases (Cherry and Williams, 2004; Lucet et al., 2006) (Figure 5F). Guided by prior studies (Kamps and Sefton, 1986), we replaced this residue with histidine in our sgRNA-resistant STK35 and PDIK1L cDNA constructs and performed competition-based proliferation assays evaluating whether these mutant cDNAs can rescue the lethality of dual targeting of STK35/PDIK1L. Despite being expressed, albeit slightly below the level of wild-type proteins, the STK35^K231H^ and PDIK1L^K37H^ behaved as null alleles in these assays, suggesting that the catalytic function of these kinase paralogs is essential in AML (Figure 5G-5H).

To further investigate whether the SCP4 and STK35/PDIK1L co-dependency in AML reflects these proteins functioning in a common genetic pathway, we performed an early time point RNA-seq analysis following acute genetic knockout of SCP4 or STK35/PDIK1L. By comparing the global transcriptional changes upon these perturbations with data obtained following knockout of other essential genes in MOLM-13, we evaluated whether SCP4 and STK35/PDIK1L were functionally linked to each other. RNA was collected on day 5 for deep sequencing analysis following lentiviral infection with SCP4 or dual STK35/PDIK1L sgRNAs, and the results were compared to datasets produced following knockout of the essential genes TAF12, MYB, SIK3, and MEF2C in this same cell line (Xu et al., 2018; Tarumoto et al., 2018). Global correlation analysis of these datasets in all pairwise combinations recapitulated the TAF12-MYB and SIK3-MEF2C connectivity described previously (Xu et al., 2018; Tarumoto et al., 2018) (Figure 5I). Importantly, the transcriptional changes observed in SCP4- and STK35/PDIK1L-deficient MOLM-13 cells were closely correlated with one another when compared to these other knockouts. The transcriptome and the co-dependency correlation suggest that the physical interaction between SCP4 and STK35/PDIK1L reflects the functioning of these signaling molecules in a common genetic pathway.

### Biochemical evidence that SCP4 functions upstream of STK35/PDIK1L

We next investigated the mechanism that links SCP4 to STK35/PDIK1L. Using western blotting and cell fractionation, we found that knockout of endogenous SCP4 led to a marked reduction in the levels of both STK35 and PDIK1L (Figure 6A and 6B). These changes in STK35/PDIK1L did not occur at the mRNA level, indicative of an alteration of the protein (Figure S6A). In contrast, a double knockout of STK35/PDIK1L did not influence the SCP4 protein (Figure S6B). The directionality of this result suggests that SCP4 functions upstream of STK35/PDIK1L to maintain the protein level of these kinases.

**Figure 6.**
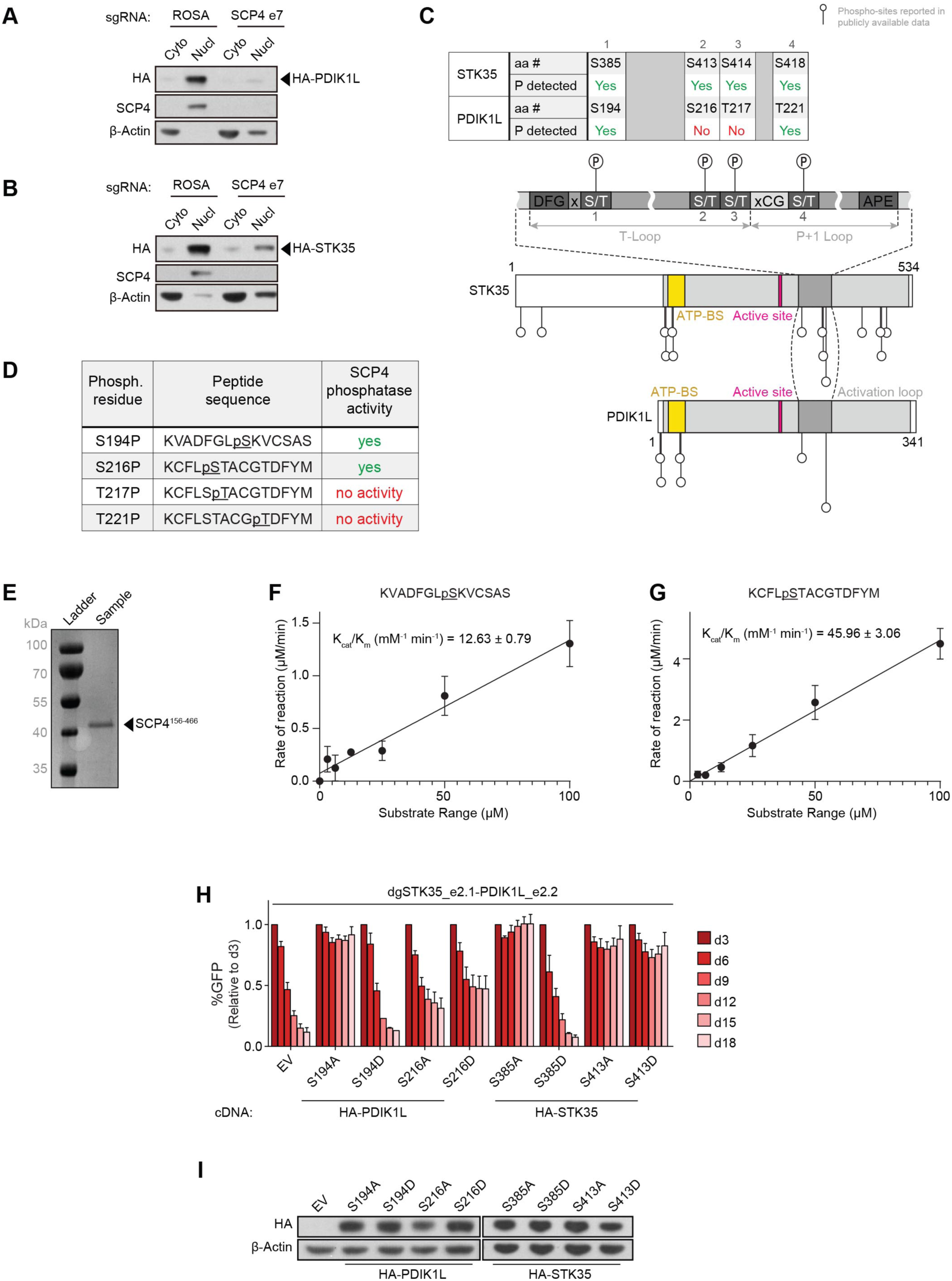
SCP4 functions upstream of STK35/PDIK1L to support AML proliferation. **(A–B)** Western blot of cytoplasm (Cyto) and nucleus (Nucl) fractions of MOLM-13 cells stably expressing HA-PDIK1L (A) or HA-STK35 (B) on day 5 post-infection with the indicated sgRNAs. Shown is a representative experiment of three independent biological replicates. **(C)** Schematics of phosphorylation sites reported in the publicly available phospho-proteomics datasets on STK35 and PDK1L relative to their domain architectures (Hornbeck et al., 2015; Ochoa et al., 2020). aa #, amino acid number; P, phosphorylation; ATP-BS, ATP-binding site. **(D)** Phosphorylated peptides assayed for SCP4 phosphatase activity *in vitro.* **(E)** Recombinant SCP4 protein purity as assessed by SDS-PAGE and Coomassie blue staining. His-SUMO-(TEV)-SCP4 was expressed in BL21 (DE3) cells and purified by affinity, anion exchange, and gel filtration chromatography. Molecular weight markers are shown for reference. **(F, G)** Phosphatase activity of SCP4 against the indicated peptides plotted for kinetic fitting. Each measurement was conducted in triplicate with standard deviations shown as error bars. **(H)** Competition-based proliferation assay in MOLM-13 cells stably expressing empty vector (EV) or the CRISPR-resistant HA-PDIK1L or HA-STK35 constructs harboring the indicated amino acid substitutions infected with the indicated sgRNAs. n = 3. **(I)** Western blot of whole-cell lysates from MOLM-13 cells stably expressing empty vector (EV) or CRISPR-resistant HA-PDIK1L or HA-STK35 constructs harboring the indicated amino acid substitutions. All bar graphs represent the mean ± SEM. All sgRNA experiments were performed in Cas9-expressing cell lines. ‘e’ refers to the exon number targeted by each sgRNA. ‘dg’ refers to the bi-cistronic vector for simultaneous targeting of STK35 and PDIK1L. The indicated sgRNAs were linked to GFP. GFP^+^ population depletion indicates loss of cell fitness caused by Cas9/sgRNA- mediated genetic mutations. The performance of negative (ROSA) and positive (PCNA) controls is summarized in Figure S6C. See also Figure S6.

We next hypothesized that the physical interaction between SCP4 and STK35/PDIK1L might allow the phosphatase to remove inhibitory phosphorylation from the kinases as a mechanism of signaling cooperativity. To investigate this, we mined publicly available proteomic databases for STK35/PDIK1L phosphorylation sites (Figure 6C) (Hornbeck et al., 2015; Ochoa et al., 2020). Among the 18 phosphorylation sites on STK35/PDIK1L identified in prior studies, we noticed several located within the regulatory activation loop of both kinases. Phosphorylation within the activation loop is a well-known mechanism to regulate kinase activity (Timm et al., 2008; Djouder et al., 2010; McCartney et al., 2016), which prompted us to investigate whether SCP4 removes phosphorylation from this segment. For this purpose, we synthesized four phosphoserine- or phosphothreonine-containing peptides representing the activation loop phosphorylated residues and optimized an expression and purification strategy for the recombinant SCP4 phosphatase domain (residue 156-466), which was well-folded and exhibited phosphatase activity using para-nitrophenyl assays (Figure 6D, 6E, and data not shown). Next, we quantified the phosphatase activity of SCP4 against each phosphopeptide using the malachite green assay to monitor phosphate release. We found that peptides corresponding to phosphoserines 194 and 216 of PDIK1L (S385 and S413 of STK35) were efficiently dephosphorylated by SCP4, whereas no activity was detected for the other two phosphopeptides (Figure 6D). The kinetic parameters of these substrates were fitted into steady-state kinetics and *kcat/Km* calculated as 12.63 ± 0.79 and 45.96± 3.06 mM-1 min-1 for phosphopeptides containing pS194 and pS216, respectively (Figure 6F and 6G).

We next performed gene complementation experiments to evaluate how alanine or phospho-mimetic aspartate substitutions of these two phosphorylated serines of STK35/PDIK1L influence AML proliferation. While all of these mutant alleles tested were expressed at comparable levels to wild-type STK35/PDIK1L, the PDIK1L^S194D^ and STK35^S385D^ proteins were completely defective in supporting AML proliferation (Figure 6H, 6I, and S6C). In contrast, alanine substitutions of these same residues were fully functional, suggesting that mutations that mimic phosphorylation inhibit the function of these kinases. Notably, the PDIK1L^S194^ and STK35^S385^ residues are located at the DFG+2 position of the activation loop for each kinase, which is reported on other kinases to form hydrogen bonds with the DFG phenylalanine to stabilize an active kinase conformation (Kornev et al., 2006; Xie et al., 2013). Collectively, these data support a model in which SCP4 removes inhibitory phosphorylation from the activation loop of STK35/PDIK1L, which may allow for kinase activation.

### SCP4-STK35/PDIK1L sustains a gene expression program of amino acid biosynthesis and transport

We next sought to understand the downstream output of this phosphatase-kinase complex that could account for its essential function in AML. From the aforementioned RNA-Seq experiments, we identified 678 genes that were significantly downregulated following both SCP4 and STK35/PDIK1L double knockout in MOLM-13 cells (Figure 7A). Ontology analysis of these genes revealed a strong enrichment for amino acid biosynthesis and transport functions (Figure 7B). For example, all the major transaminases that catalyze reversible interconversion of glutamate into other amino acids were downregulated as well as many of the solute carriers that import amino acids into the cell (Figure 7C). To evaluate whether these mRNA changes lead to corresponding changes in cellular amino acid levels, we performed MS to measure metabolites following SCP4 or STK35/PDIK1L inactivation in MOLM-13 cells. While the level of several metabolites remained unchanged in these knockout cells, many amino acids were present at reduced levels in cells deficient for either SCP4 or STK35/PDIK1L (Figure 7D). One of the most deficient amino acids was proline, which could be related to the decrease in *PYCR1* expression, encoding the rate-limiting enzyme for proline biosynthesis (Xiao et al., 2020) (Figure S7A). Since amino acid biosynthesis and transport are known to be deregulated to support the rapid growth of AML cells (Bertuccio et al., 2017; Willems et al., 2013; Bjelosevic et al., 2021; Kreitz et al., 2019), these results lead us to propose that SCP4-STK35/PDIK1L signaling supports the aberrant metabolic state of this malignancy (Figure 7E).

**Figure 7.**
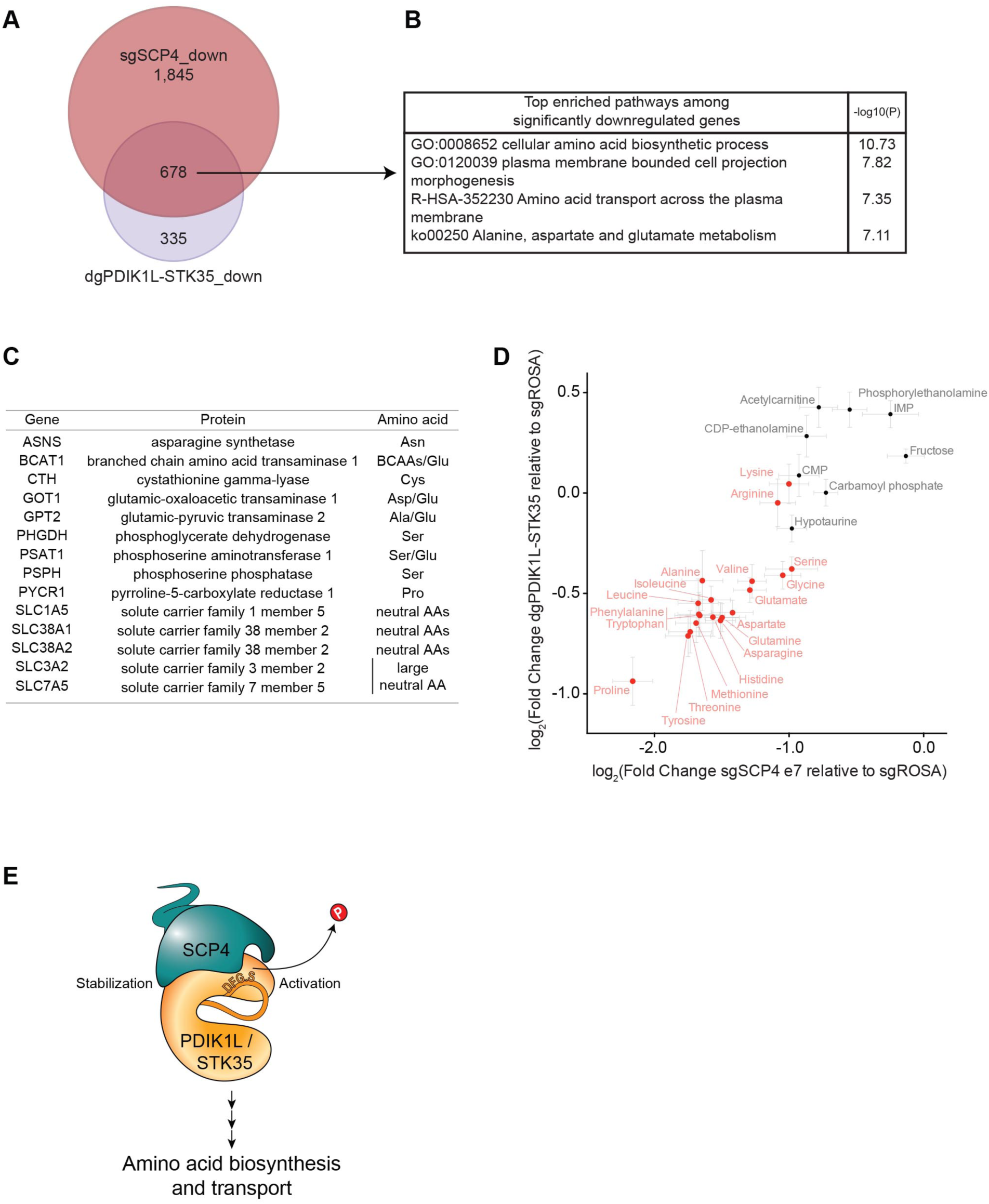
SCP4-STK35/PDIK1L complex is needed to sustain amino acid levels in AML. **(A)** Venn diagram depicting the overlap between statistically significant downregulated genes in MOLM-13 cells upon SCP4 knockout and STK35/PDIK1L double knockout. DeSeq2 (n = 4). **(B)** Ontology analysis of overlapping statistically significant downregulated genes in MOLM-13 cells upon both SCP4 knockout and STK35/PDIK1L double knockout. **(C)** Selected statistically significant downregulated genes in MOLM-13 cells upon both SCP4 knockout and STK35/PDIK1L double knockout and the amino acids they are involved in biosynthesis or transport of. **(D)** The correlation between log2 fold changes in the levels of selected metabolites upon SCP4 knockout and STK35/PDIK1L double knockout relative to negative control as measured by the MS analysis. Every dot represents the mean ± SEM (n = 6). Amino acids are in red, few unchanged metabolites in black for reference. Shown is a representative experiment of three independent biological replicates. **(E)** Model. See also Figure S7.

## DISCUSSION

The main conclusion of this study is that the nuclear phosphatase SCP4 is an acquired dependency in AML. Using biochemical and genetic approaches, we found that SCP4 executes this function by associating with the kinase paralogs STK35 and PDIK1L. Our experiments implicate both the catalytic phosphatase and kinase activities within this complex as functioning in the same AML maintenance pathway. We reveal that one downstream output of this signaling complex is to maintain the expression of amino acid biosynthetic enzymes and transporters. These findings lead us to speculate that the SCP4-STK35/PDIK1L complex is required in AML to support the aberrant metabolic state in this disease. In addition to its therapeutic implications, our study illustrates the capability of high-throughput CRISPR-Cas9 genetic screens in revealing novel signaling pathways in cancer.

While several phosphatases are known to regulate kinase activities by removal of phosphorylation on the activation loop (Takekawa et al., 1998; Salmeen et al., 2000; Cheng et al., 2000; Hayward et al., 2019), relatively few examples exist of docking of phosphatase and kinase proteins to form a stable complex. One prominent example of such an interaction is the phosphatase KAP and its interaction with its kinase substrate CDK2 (Hannon et al., 1994; Poon and Hunter, 1995). X-ray crystallography revealed that the binding interface between these two proteins involves extensive interactions remote from the active sites of both proteins, which position CDK2 for removal of activating phosphorylation of its activation loop (Song et al., 2001). As another example, the phosphatase MKP3 stably associates with its kinase substrate ERK2 via a docking surface distinct from the catalytic active sites of each enzyme (Zhang et al., 2003; Liu et al., 2006). In each of these examples, the docking interaction leads to allosteric modulation of catalytic kinase and/or phosphatase activity. We speculate that a similar docking interaction and potential for allostery exists between SCP4 and STK35/PDIK1L, which may lead to stabilization of the kinase substrate and positioning of the kinase substrates for de-phosphorylation.

One paradoxical result in our study is the strength of the association between the catalytic domains of SCP4 and STK35/PDIK1L, yet this interaction is weakened by a single point mutation of the SCP4 active site. Since the interaction of enzymes with their substrates are often transient (Fahs et al., 2016), it is unlikely that the strong binding interaction we detect between SCP4 and STK35/PDIK1L is entirely due to an active site interaction with a phosphorylated substrate. While our experiments indicate that the activation loop of STK35/PDIK1L is a substrate of SCP4, we speculate that the binding interaction between SCP4 and STK35/PDIK1L includes a docking mechanism involving non-catalytic surfaces of these two proteins. One possibility is that the SCP4^D293A^ protein becomes locked in an inactive conformation, which simultaneously impairs both catalysis and docking with STK35/PDIK1L. Future structural studies of the SCP4-STK35/PDIK1L complex coupled with deep mutagenesis could shed further light on this specific binding surface that links these two catalytic domains.

Our biochemical assays and mutational analyses implicate a functionally important activation loop serine of STK35 (S385) and PDIK1L (S194) as substrates of SCP4. These serines lie at the +2 position relative to the DFG motif, a region of the activation loop that binds to a magnesium ion that interacts directly with ATP. Substantial evidence suggests that the DFG motif can adopt distinct open or closed conformations to regulate kinase activity (Kornev et al., 2006; Xie et al., 2020). While activation loop phosphorylation is known to promote kinase activity from several downstream locations, we hypothesize that incorporation of phosphorylation at the DFG+2 serine residue of STK35/PDIK1L locks the activation loop in inactive conformation that is incompatible with the precise ATP-magnesium coordination. This is consistent with the requirement for the 3-turn formation between the DFG phenylalanine and the DFG+2 residue, reported for other active kinases (Kornev et al., 2006; Xie et al., 2013).

The neonatal lethality phenotype of SCP4^-/-^ mice can be rescued by injection of glucose, which suggests a fundamental role of SCP4 in regulating metabolic pathways *in vivo* (Cao et al., 2018). In this prior study, SCP4 was found to regulate FoxO transcription factors in hepatocytes which in turn influences the expression of critical genes in the gluconeogenesis pathway (e.g., *PCK1* and *G6PC*). In AML, we find that SCP4 regulates instead the expression of a distinct group of metabolic genes that control amino acid biosynthesis and transport. Thus, it appears that distinct transcriptional outputs exist downstream of SCP4 in different cell types. A key question raised by our study is whether a specific transcription factor might be regulated by SCP4 or by STK35/PDIK1L to influence amino acid biosynthesis/transport genes. Interestingly, we found that many SCP4-regulated genes involved in metabolism are also target genes of the stress-induced transcription factor ATF4 (Harding et al., 2003). In the future, it would be interesting to determine whether SCP4 or STK35/PDIK1L directly modify the phosphorylation state of this protein to influence metabolism.

Several HAD family phosphatases have been successfully targeted using small-molecule inhibitors (Freschauf et al., 2009; Krueger et al., 2014; Wang et al., 2016). Thus, the acquired dependency of AML cells on SCP4 presents an attractive target for cancer therapy. In addition, the kinase activity of STK35/PDIK1L also might be targetable using small molecules that compete with ATP binding to these targets. Our study provides justification for the development of such tool compounds targeting this phospho-catalytic signaling complex, which would be expected to suppress AML cells while having minimal effects on normal hematopoiesis.

## AUTHOR’S CONTRIBUTIONS

S.A.P. and C.R.V. designed experiments and analyzed results, S.A.P. performed and analyzed experiments. R.Y.M., S.I., and Y.J.Z. purified and measured phosphatase activity of SCP4. B.L. performed and analyzed SCP4 CRISPR exon scan. S.A.P., R.F. and Y.Y. performed editing in human CD34^+^ cells. S.A.P., O.K., and O.E.D. analyzed RNA-Seq data. O.K. provided bi-cistronic vector for double knockout. Z.Y. performed and analyzed cell cycle arrest and apoptosis experiments. Y.W. performed and analyzed mouse xenograft experiments. L.A.B. performed several competition assays. S.A.P. and C.R.V. wrote the manuscript.

## ACKNOWLEDGMENTS

We are thankful to the members of the Vakoc laboratory and Weiss and Zhang laboratories for insightful discussions and helpful suggestions throughout this study. We thank CSHL Functional Genomics Core Facility, specifically Kenneth Chang for helpful discussions; CSHL Next Generation Sequencing Core Facility for deep sequencing; and CSHL Bioinformatics Shared Resource for providing the Galaxy data analysis platform. We thank the Proteomics & Metabolomics Facility at the Center for Biotechnology/University of Nebraska-Lincoln for the mass spectrometry protein identification from affinity purification experiments, specifically Mike Naldrett and Sophie Alvarez. We thank CSHL Mass Spectrometry Shared Resource supported by the Cancer Center Support Grant 5P30CA045508 for the metabolites mass spectrometry measurements, specifically Sofia Costa. We would also like to thank Scott Lyons and Michael Lukey for helpful discussions during the manuscript preparation. This work was supported by Cold Spring Harbor Laboratory NCI Cancer Center Support grant 5P30CA045508. Additional funding was provided to C.R.V. by the Pershing Square Sohn Cancer Research Alliance, National Institutes of Health grants R01 CA174793 and P01 CA013106, and a Leukemia & Lymphoma Society Scholar Award.

## MATERIALS AND METHODS

### Cell lines

Human cell lines MOLM-13, NOMO-1, MV4-11, ML-2, HEL, SET-2, THP-1, U937 (acute myeloid leukemia, AML); K562 (chronic myelogenous leukemia, CML); PANC-1, MIAPaCa-2, SUIT-2, AsPC-1 (pancreatic ductal adenocarcinoma, PDAC); RH4, RH30 (rhabdomyosarcoma, RMS); NCI-H1048 (small cell lung cancer, SCLC); and murine cell line RN2 (MLL-AF9/NRas^G12D^ AML) (Zuber et al., 2011) were cultured in RPMI-1640 supplemented with 10% fetal bovine serum (FBS). Human cell line SEM (AML) was cultured in IMDM with 10% FBS. Human cell line OCI-AML3 (AML) was cultured in Alpha MEM with 20% FBS. Transformed human HSPC MLL-AF9 (MA9) cells (Mulloy et al., 2008; Wei et al., 2008) derivatives MA9-NRAS^G12D^ and MA9-FLT3^ITD^ (AML) were cultured in IMDM with 20% FBS. Human cell line Kasumi-1 (AML) was cultured in RPMI with 20% FBS. Human cell lines RD (RMS), A549 (lung adenocarcinoma), and HEK293T were cultured in DMEM with 10% FBS. Murine cell line NIH3T3 was cultured in DMEM with 10% bovine calf serum (BCS). 1% penicillin/streptomycin was added to all media. All cell lines were cultured at 37°C with 5% CO_2_ and were regularly tested mycoplasma negative.

### Plasmid construction and sgRNA cloning

The sgRNA lentiviral expression vector with optimized sgRNA scaffold backbone (LRG2.1, Addgene plasmid # 108098) was derived from a lentiviral U6-sgRNA-EFS-GFP expression vector (LRG, Addgene plasmid # 65656) by replacing the original wild-type sgRNA scaffold with our optimized version (Tarumoto et al., 2018). LRCherry2.1 was produced from LRG2.1 by replacing GFP with mCherry CDS. LRG2.1-Blast was generated by the introduction of the blasticidin resistance gene after GFP CDS with a P2A linker. In this study, LRG2.1, LRG2.1-Blast, or LRCherry2.1 were used for introducing sgRNA into human cell lines, while the original LRG vector was used for sgRNA expression in murine cell lines. All the sgRNAs were cloned into the LRx2.1-derivatives or LRG vectors using a BsmBI restriction site. Sequences of all sgRNAs used in this study are provided in the supplemental Table S3 (Table S3. sgRNAs sequences, CRISPR-resistant constructs, and primers. Related to all Figures).

LentiV_Neo vector was derived from LentiV_Cas9_puro vector (Addgene plasmid # 108100) by removing Cas9 and replacing a puromycin resistance gene with a neomycin resistance gene. *CTDSPL2* cDNA (Horizon Discovery, Clone ID: 5744745), partial *STK35* cDNA (Horizon Discovery, Clone ID: 9021751), and *PDIK1L* cDNA (Horizon Discovery, Clone ID: 4828997) were cloned into LentiV_Neo vector using In-Fusion cloning system (Takara Bio; Cat. No. 121416). The N-terminus of STK35 cDNA (4–296 bp from translation start site) was added to obtain STK35 cDNA corresponding to NM_080836. The N-terminal FLAG and HA-tags were added to cDNAs in-frame. For the generation of all the mutants, the base substitutions were introduced into the cDNA via PCR mutagenesis. The list of all the mutants used in this study can be found in the Supplemental Table S3 (Table S3. sgRNAs sequences, CRISPR-resistant constructs, and primers. Related to all Figures).

### Construction of sgRNA libraries

All the genes contained within the gene group “Phosphatases” reported by the HUGO Gene Nomenclature Committee (HGNC) were selected for designing the phosphatase domain-focused sgRNA library. The phosphatase domain information was retrieved from the NCBI Conserved Domains Database. Six to thirty independent sgRNAs were designed against exons encoding phosphatase domains. For the CRISPR exon-scan library 85 sgRNAs were selected against every exon of SCP4. All sgRNAs were designed using https://mojica.cshl.edu and filtered for the minimal predicted off-target effects (Hsu et al., 2013).

The domain targeting and positive/negative control sgRNAs oligonucleotides were synthesized in a pooled format on an array platform (Twist Bioscience) and then amplified by PCR, using Phusion® Hot Start Flex DNA Polymerase (NEB; Cat. No. M0535). The PCR products were cloned into the BsmBI-digested lentiviral vector LRG2.1 (Addgene plasmid # 108098) using a Gibson Assembly^®^ Cloning Kit (New England BioLabs; Cat. No. E2611). The library was produced in MegaX DH10B T1R Electrocomp Cells (Invitrogen; Cat. No. C640003).

### Lentivirus production and infection

The lentivirus was packed in HEK293T cells by transfecting lentiviral expression vector plasmids together with the lentiviral packaging plasmid (psPAX2, Addgene plasmid # 12260) and the envelope protein expressing plasmid (VSV-G) using polyethylenimine (PEI 25K™; Polysciences; Cat. No. 23966) transfection reagent. HEK293T cells were transfected when ∼80–90% confluent. For the pooled sgRNA phosphatase library lentivirus production, five 10 cm plates of HEK293T were used. For one 10 cm plate of HEK293T cells, 5 μg of plasmid DNA, 5 μg of VSV-G, 7.5 μg psPAX2, and 64 μL of 1 mg/mL PEI were mixed with 1 mL of Opti-MEM®, incubated at room temperature for 20 minutes, and then added to the cells. The media was changed to 5 mL of fresh media 6–8 hours post-transfection. The lentivirus-containing media was collected at 24, 48, and 72 hours post-transfection, pooled together and filtered through a 0.45 μM non-pyrogenic filter. For lentivirus infection, target cells were mixed with the virus (volume empirically determined to result in the desired infection rate) and 4 μg/ml polybrene and then centrifuged at 600 x g for 30– 45 minutes at room temperature. For adherent cell lines, media was changed at 24 hours post-infection. If selection was required for stable cell line establishment, corresponding antibiotics (1 μg/mL puromycin, 1 μg/mL blasticidin, 1 mg/mL G418) were added 24 hours post-infection.

### Pooled negative-selection CRISPR screening and data analysis

Cas9-expressing stable cell lines were produced by infection with the lentivirus encoding human codon-optimized *Streptococcus pyogenes* Cas9 protein (LentiV_Cas9_puro, Addgene plasmid # 108100) and subsequent selection with 1 μg/mL puromycin. Lentivirus of pooled sgRNA phosphatase library titer was estimated by mixing the cells with the serially diluted virus and measuring GFP% on day three post-infection using a Guava® easyCyte™ Flow Cytometer (Merck Millipore). The cells were then infected with the volume of virus estimated to result in a 30–40% infection rate to increase the probability of a single sgRNA introduction event per cell. The number of cells was kept 1000 times more than the sgRNA number in the library to maintain the representation of sgRNAs during the screen. Cells were harvested at initial (day three post-infection) and final (10 or more population doublings after the initial) time points. Genomic DNA was extracted using the QIAamp DNA Mini Kit (QIAGEN; Cat. No. 51304).

Sequencing libraries were prepared as described previously (Shi et al., 2015). Briefly, genomic DNA fragments (∼200 bp) containing sgRNAs were amplified by PCR, using Phusion® Hot Start Flex DNA Polymerase (NEB; Cat. No. M0535). The PCR products were end-repaired with T4 DNA polymerase (NEB; Cat. No. B02025), DNA Polymerase I, Large (Klenow) fragment (NEB; Cat. No. M0210L), and T4 polynucleotide kinase (NEB; Cat. No. M0201L). The 3’ adenine overhangs were added to the blunt-end DNA fragments by Klenow Fragment (3’-5’ exo; NEB; Cat. No. M0212L). The DNA fragments were then ligated with diversity-increased custom barcodes (Shi et al., 2015), using Quick Ligation Kit (NEB; Cat. No. M2200L). The ligated DNA was PCR amplified with primers containing Illumina paired-end sequencing adaptors, using Phusion Flash High-Fidelity PCR Master Mix (Thermo Fisher; Cat. No. F548S). The final libraries were quantified using Bioanalyzer Agilent DNA 1000 (Agilent 5067-1504) and were pooled together in an equimolar ratio for paired-end sequencing using MiSeq platform (Illumina) with MiSeq Reagent Kit V3 150-cycle (Illumina).

The sgRNA sequences were mapped to the reference sgRNA library to discard any mismatched sgRNA sequences. The read counts were calculated for each individual sgRNA. The following analysis was performed with a custom Python script: sgRNAs with read counts less than 50 in the initial time point were discarded; the total read counts were normalized between samples; the average log_2_ fold-change in the reads corresponding to sgRNAs targeting a given gene (CRISPR score) was calculated, as described (Wang et al., 2015). AML-specific dependency was determined by subtracting the average of CRISPR scores in non-AML cell lines from the average of CRISPR scores in AML cell lines, and that score was ranked in ascending order. The phosphatase CRISPR screening data is provided in the Supplemental Table S1 (Table S1. Average log_2_ FC in phosphatase domain-focused screen. Related to Figure 1 and S1). The SCP4 CRISPR exon scanning screening data is provided in the Supplemental Table S2 (Table S2. Average log_2_ fold-change in SCP4 CRISPR exon scanning screen in MOLM-13. Related to Figure 3 and S3).

### Competition-based cell proliferation assays

Cas9-expressing cell lines were infected with lentivirus encoding sgRNAs linked with either GFP or mCherry reporters at the infection rate of ∼20–60% as described above. Percentage of GFP- or mCherry-positive cells was measured every three days from day 3 to day 18 post-infection using Guava® easyCyte™ Flow Cytometer (Merck Millipore). The percentage of fluorescent cells in the population on each day was divided by their percentage on day three to calculate fold-change corresponding to the effect of a given sgRNA on cell proliferation.

### Western Blot

For knock-out experiments, cells were harvested on day 5 post-infection with sgRNA.

For whole-cell lysates, a precise cell number was calculated before sample preparation for equal loading. Cells were washed 1X with ice-called PBS. Cell pellets were then resuspended in PBS and 2X Laemmli Sample Buffer (Bio-Rad, Cat. No. 1610737) containing 2-mercaptoethanol, boiled for 30 minutes, and cleared by centrifugation at room temperature at 13,000 rpm.

For fractionation experiments, cell fractions were prepared as described below. The exact protein concentrations in the fractions were measured with the Pierce™ BCA Protein Assay Kit (Thermo Fischer, Cat. No. 23225). The cell fractionation samples were boiled for 5–10 minutes with a 2X Laemmli Sample Buffer containing 2-mercaptoethanol.

The samples were separated by SDS-PAGE (NuPAGE 4-12% Bis-Tris Protein Gels, Thermofisher), followed by transfer to nitrocellulose membrane via wet transfer at 90 V for 2–2.5 hours. Membranes were then blocked for 1 hour with 5% non-fat milk in TBST and incubated with primary antibodies overnight. On the next day, the membranes were washed 3X with TBST followed by incubation with HRP-conjugated secondary antibodies (Rabbit Cytiva/Amersham NA934; Mouse Agilent/Dako P026002-2) for one hour at room temperature. After 3X washes with TBST, an enhanced chemiluminescence (ECL) solution containing HRP substrate was added to the membranes, and the signal was visualized using the darkroom development techniques for chemiluminescence. Antibodies used in this study included SCP4 (*CTDSPL2*) (CST, # 6932, 1:500), FLAG (Sigma Aldrich, F1804, 1:10,000), HA-HRP (Sigma Aldrich, clone 3F10, 1:10,000), H3K4me3 (Sigma Aldrich, 07-473, 1:1,000), β-Actin-HRP (Sigma A3854-200UL; 1:50,000).

### Cell cycle arrest and apoptosis analysis

Cell cycle analysis was performed using the BrdU Flow Kit according to the manufacturer protocol (BD, FITC BrdU Flow Kit; Cat. No. 559619), with cells pulsed with BrdU for 1 hour at 37°C. Annexin V apoptosis staining was performed using the Apoptosis Detection Kit according to the manufacturer protocol (BD, FITC Annexin V Apoptosis Detection Kit; Cat. No. 556547). Cells were co-stained with 4′,6-diamidino-2-phenylindole (DAPI) to measure DNA content and analyzed with a BD LSRFortessa flow cytometer (BD Biosciences) and FlowJo software (TreeStar).

### Editing and differentiation of human peripheral blood CD34^+^ cells

Circulating G-CSF-mobilized human CD34^+^ cells were obtained from two de-identified healthy donors (KeyBiologics). Enrichment of CD34^+^ cells was performed by immunomagnetic bead selection using an CliniMACS instrument (Miltenyi Biotec). After enrichment, the CD34^+^ cell fraction was 95% of the total, with <0.2% CD3^+^ cells and <2% CD19^+^ cells.

Purified recombinant Cas9 protein was obtained from Berkeley Macrolabs. Two sgRNAs against SCP4 and one sgRNA against MYC were synthesized by TriLink BioTechnologies. The sgRNAs included 2′-O-methyl 3′-phosphorothioate (MS) modifications at 3 terminal nucleotides at both the 5′ and 3′ ends (Hendel et al., 2015).

Editing in human CD34^+^ cells was performed as described before (Metais et al., 2019). Briefly, cryopreserved CD34^+^ cells were thawed and pre-stimulated for 48 hours in the maintenance medium (Table S4). The cells were then washed 1X in PBS, resuspended in T buffer included in kit (Thermo Fisher Scientific, MPK10025), mixed with RNP (described below), and electroporated with 1600 V, 3 pulses of 10 ms, using a Neon Transfection System (Thermo Fisher Scientific, Cat. No. MPK5000). As a control, cells were not electroporated or electroporated with Cas9 and Non-targeting gRNA.

Ribonucleoprotein complexes (RNPs) were formed by incubating 160 pmol of Cas9 with 320 pmol sgRNA in 50 μL of 10 mM HEPES (Thermo Fisher Scientific, catalog # 15630080), 150 mM NaCl (Thermo Fisher Scientific, catalog # 9759) for 35 minutes. 1 million CD34^+^ cells resuspended in 50 μL of T buffer were mixed with RNP complex in a final volume of 100 μL.

After electroporation, cells were immediately added to the media with cytokines required for differentiation into erythroid, myeloid, or megakaryocyte lineages or to methylcellulose for the colony forming cell (CFC) assay (see below). Erythroid, myeloid, and megakaryocyte maturation were monitored by immune-flow cytometry for the appropriate cell surface markers (Table S4) on days 2, 5, 8, and 16 post-electroporation (final day for differentiation) using BD LSRFortessa Dual™ Flow Cytometer. The genomic DNA from each of the lineages was collected in parallel to the immune-flow cytometry, and Western blot samples were prepared on days 5 and 8.

The on-target indel frequencies were quantified using QuantStudio™ 12K Flex Real-time PCR System (Thermo Fisher Scientific) with Power SYBR Green Master Mix (Thermo Fisher). Two primer pairs were developed for each targeted region using Primer3 (Untergasser et al., 2012). One primer in each pair was chosen to span the Cas9 cleavage site 3-nt upstream of the PAM site, and the second primer was selected at a 100–150 bp distance. Primer sequences are provided in Table S4. Quantitative PCR was performed on genomic DNA from gene-edited cells or control cells. Based on the consistency of amplification, either chr10 or untargeted region of CTDSPL2 was used as an endogenous control for calculating ΔCT values. The percentage of editing in CD34^+^ cells electroporated with sgRNAs vs. Cas9 alone was approximated as 1 - 2^(-ΔΔCT). For the myeloid lineage, Surveyor nuclease assay was performed with primers in Table S4, according to the manufacturer protocol (Integrated DNA Technologies, Cat. No. 706020) (Table S4. sgRNAs; media and cytokines; and antibodies used in flow cytometry panels in CD34^+^ experiments. Related to Figure 2 and S2).

For the CFC assay, ∼800 CD34^+^ gene-edited and control cells were mixed with 1.0 mL of MethoCult™ SF H4636 methylcellulose (Stemcell Technologies, Cat. No. 04636) containing recombinant cytokines for human embryonic stem (ES) cell-derived hematopoietic progenitor cells. The cultures were incubated in 35-mm tissue culture dishes. After 14 days, single colonies were classified and quantified.

All the reagents used in these experiments can be found in the Supplemental Table S4 (Table S4. sgRNAs; media and cytokines; and antibodies used in flow cytometry panels in CD34^+^ experiments. Related to Figure 2 and S2).

### *In vivo* transplantation of MOLM-13 cells into NSG mice

All animal procedures and studies were approved by the Cold Spring Harbor Laboratory Animal Care and Use Committee in accordance with IACUC. First, MOLM-13/Cas9^+^ cell line stably expressing luciferase was established via lentiviral infection with a Lenti-luciferase-P2A-Neo (Addgene # 105621) vector followed by G418 (1 mg/mL) selection. These cells were infected with lentivirus encoding GFP-linked sgRNAs targeting either *ROSA26* locus (negative control) or *CTDSPL2* (SCP4). On day 3 post-infection, the infection rate was checked by the percentage of GFP positive cells, and all samples had over 90% infection rate. 0.5 million cells were transplanted into sublethally (2.5Gy) irradiated NSG mice (Jax 005557) through tail vein injection. Mice were imaged with IVIS Spectrum system (Caliper Life Sciences) on days 12 and 15 post-injection for visualizing the disease progression.

### Sub-cellular fractionation and Immunoprecipitation in MOLM-13 cells

Subcellular protein fractionation was performed either by using Subcellular Protein Fractionation Kit for Cultured Cells (Thermo Scientific; Cat. no. 78840) according to the manufacturer protocol or following the protocol below.

For subsequent mass spectrometry (MS) analysis, ∼200 million cells were collected and the protocol below was used. For immunoprecipitation (IP) coupled with Western blot (WB), 40–60 million cells were collected and the protocol below was used.

The cells were washed 2X with ice-cold PBS, then resuspended in 5X Packed Cell Volume (PCV) of Hypotonic cell lysis buffer (10 mM HEPES pH 7.9; 1.5 mM MgCl2; 10 mM KCl; 1 mM DTT; supplemented with proteinase inhibitors). Note: SCP4 might be a target of myeloid proteases in MOLM-13 cells (data not shown) (Zhong et al., 2018). Therefore, we found that the best protein preservation was achieved when avoiding the use of detergents in the protocol and thoroughly washing the nuclei pellet from any traces of the cytoplasm. After 15 minutes incubation at 4°C with rotation, the cells were pelleted and resuspended in 2X PCV of Hypotonic cell lysis buffer. The cell walls were disrupted using a glass tissue homogenizer on ice with 10 up-and-down strokes using a type B pestle. The disrupted cells in suspension were pelleted for 20 minutes at 11,000 x g at 4°C. The supernatant was saved as the *cytoplasmic fraction*. The pellet was thoroughly washed 4X with Hypotonic cell lysis buffer. The crude nuclei pellet was resuspended in 2/3X PCV of Extraction Buffer (20 mM HEPES pH 7.9; 1.5 mM MgCl2; 420 mM NaCl; 25% Glycerol; 1 mM DTT; 1:1000 benzonase (Sigma E1014-25KU); supplemented with proteinase inhibitors). The nuclear lysates were incubated at room temperature for 1 hour on the rotor and centrifuged at 4°C for 10 minutes at 21,000 x g. The supernatant was diluted to 150 mM NaCl using No-salt dilution buffer (20 mM HEPES pH 7.9; 1.5 mM MgCl2; 0.31 mM EDTA; 1 mM DTT; supplemented with proteinase inhibitors) and saved as the *nuclear fraction*. The exact protein concentrations in the fractions were measured with the Pierce™ BCA Protein Assay Kit (Thermo Fischer, Cat. No. 23225).

For immunoprecipitation, NP-40 was added up to 0.25%, and the nuclear fractions were incubated with anti-FLAG M2 agarose (Sigma A2220-10ML) or anti-HA agarose (Sigma E6779-1ML) or Pierce Anti-HA Magnetic Beads (Thermofisher 88837) at 4°C overnight. Next day, the beads were washed 5X with Wash buffer (20 mM HEPES pH 7.9; 150 mM NaCl; 0.2 mM EDTA; 0.25% NP-40; 1 mM DTT; supplemented with proteinase inhibitors). For WB the beads were boiled for 10 minutes in 1X Laemmli Sample Buffer (Bio-Rad, Cat. No. 1610737) containing 2- mercaptoethanol. For MS, the beads were send on ice to the Proteomics and Metabolomics Facility Center for Biotechnology at the University of Nebraska–Lincoln.

### Protein identification by mass spectrometry (MS)

Bead samples were made up in 60μL 1x non-reducing LDS sample buffer and incubated at 95°C for 10 min prior to loading 50μL of sample onto a Bolt™ 12% Bis-Tris-Plus gel and running briefly into the top of the gel. The gel was fixed and stained with colloidal coomassie blue G250 stain. Gel containing the proteins was reduced and alkylated, then washed to remove SDS and stain before digestion with trypsin (500ng) overnight at 37°C. Peptides were extracted from the gel pieces, dried down, and samples were re-dissolved in 30μL, 2.5% acetonitrile, 0.1% formic acid. 5μL of each digest was run by nanoLC-MS/MS using a 2h gradient on a 0.075 mm x 250 mm CSH C18 column feeding into a Q-Exactive HF mass spectrometer.

All MS/MS samples were analyzed using Mascot (Matrix Science, London, UK; version 2.6.2). Mascot was set up to search the cRAP_20150130.fasta (123 entries); uniprot-human_20191024 database (selected for Homo sapiens, unknown version, 74,034 entries) assuming the digestion enzyme trypsin. Mascot was searched with a fragment ion mass tolerance of 0.060 Da and a parent ion tolerance of 10.0 PPM. Deamidated of asparagine and glutamine, oxidation of methionine, carbamidomethyl of cysteine, phospho of serine, threonine and tyrosine were specified in Mascot as variable modifications.

Scaffold (version Scaffold_4.8.9, Proteome Software Inc., Portland, OR) was used to validate MS/MS based peptide and protein identifications. Peptide identifications were accepted if they could be established at greater than 80.0% probability by the Peptide Prophet algorithm (Keller et al., 2002) with Scaffold delta-mass correction. Protein identifications were accepted if they could be established at greater than 99.0% probability and contained at least 2 identified peptides. Protein probabilities were assigned by the Protein Prophet algorithm (Nesvizhskii et al., 2003). Proteins that contained similar peptides and could not be differentiated based on MS/MS analysis alone were grouped to satisfy the principles of parsimony. Proteins sharing significant peptide evidence were grouped into clusters.

### Co-IP in HEK 293T cells

One 80% confluent 10-cm plate of 293T cells was transfected with 10 μg HA-PDIK1L and 10 μg of either Empty vector, or FLAG-SCP4, or FLAG-Δ235aa-SCP4, or FLAG-SCP4^D293A^. Total 20 μg DNA was mixed with 60 μL of 1 mg/mL PEI and 1 mL of Opti-MEM®, incubated at room temperature for 15 minutes, and then added to the cells. 24 hours post-transfection the whole-cell lysates were prepared as follows. Collected cells by incubation for 5 minutes with 2 mL Trypsin, then neutralized with 6 mL DMEM media. Washed cells 2X with ice-cold PBS. Resuspended cells in 500 μL of ice-cold Cell Lysis Buffer (CST, #9803). Incubated on ice for 30 minutes. Centrifuged at 4°C for 10 minutes at 14,000 x g. The exact protein concentrations in the fractions were measured with the Pierce™ BCA Protein Assay Kit (Thermo Fischer, Cat. No. 23225). Whole-cell lysates were incubated with anti-FLAG M2 agarose (Sigma A2220-10ML) or anti-HA agarose (Sigma E6779-1ML). Next day, the beads were washed 5X with the Cell Lysis Buffer. Afterwards, the beads were boiled for 10 minutes in 1X Laemmli Sample Buffer (Bio-Rad, Cat. No. 1610737) containing 2-mercaptoethanol, followed by Western blot analysis.

### RNA-Seq

For knock-out experiments, cells were harvested on day 5 post-infection with sgRNA. RNA was purified with the guanidinium thiocyanate-phenol-chloroform extraction method. Briefly, cells were washed 1X with PBS and lysed with 1 mL of TRIzol (Thermo Scientific; Cat. No. 15596018). The samples were vigorously mixed with 200 µL chloroform and incubated for 3 minutes at room temperature, followed by centrifugation at 10,000 x g for 15 minutes at 4 °C. The aqueous phase containing RNA was added to the equal volume of isopropanol, and RNA was precipitated after incubation of 10 minutes at room temperature. Precipitated RNA was then pelleted by centrifugation at 10,000 x g for 10 minutes at 4 °C and washed 1X with 1 mL of 75% ethanol. After air-drying for 10 minutes, RNA was resuspended in 20 µL of DEPC-treated (RNase-free) water.

RNA-seq libraries were prepared using TruSeq sample prep kit v2 (Illumina) according to the manufacturer protocol. Briefly, polyA RNA was enriched with oligo-dT beads and fragmented enzymatically. The cleaved RNA fragments were reverse transcribed into first strand cDNA using SuperScript II reverse transcriptase and random primers. The RNA template was then removed, and a replacement strand to generate double-stranded (ds) cDNA was synthesized. The ds cDNA was purified with AMPure XP beads, end-repaired, 3’-adenylated, and ligated with indexing adapters, preparing them for hybridization onto a flow cell. The DNA fragments that have adapter molecules on both ends were selectively enriched by PCR-amplification and purified with AMPure XP beads. RNA-seq libraries were pooled together in equimolar concentrations and analyzed by single-end sequencing using NextSeq (Illumina). Four independent biological replicates for control samples, samples from cells with SCP4 knock-out, and cells with STK35-PDIK1L double knock-out were sequenced in the same flow cell.

### RNA-Seq data analysis

Sequencing reads were mapped into reference human genome hg19 using STAR v.2.5.2b-1 (Dobin et al., 2013). The mapped reads were assigned to genes using HTSeq-Count v.0.6.1p1 (Anders et al., 2015). The differential expression gene analysis was performed using DESeq2, with four replicates for each sample (Love et al., 2014). Pesudo-alignment to the reference human transcriptome (gencode v.35), assignment of reads to genes, and TPM calculation were all performed using Salmon (v.1.0) (Patro et al., 2017). For comparison with other RNA-Seq datasets, raw RNA-Seq files from GSE109491 and GSE104308 datasets corresponding to the available replicates of negative controls, sgTAF12, sgMYB, sgSIK3, and sgMEF2C were downloaded. All the raw datasets, including ours, were re-aligned (pseudo-alignment) using Kallisto to hg38, bootstrap 100 (Bray et al., 2016).The differential expression gene analysis was performed using DESeq2, with four replicates for each sample (Love et al., 2014). Pearson’s product-moment correlations between global log_2_ fold changes observed upon different knockouts in MOLM-13 cells were calculated using *cor()* function in R with default parameters.

### Protein Expression and Purification

The SCP4 phosphatase domain (encoding residues 156–466) was subcloned into a pET28a (Novagene) derivative vector encoding a 6xHis-tag followed by a SUMO-tag and a TEV protease site. BL21 (DE3) cells expressing SCP4 were grown in one-liter cultures at 37°C in Luria-Bertani (LB) broth (Thermo) containing 50µg/ml kanamycin. Once the cultures reached an OD_600_ value of 0.6–0.8, the protein expression was induced with 0.2 M isopropyl-β-D-thiogalactopyranoside (IPTG), and the cultures were grown an additional 16 hours at 18°C. The cells were pelleted and resuspended in lysis buffer (50 mM Tris-HCl pH 8.0, 500 mM NaCl, 15 mM imidazole, 10% glycerol, 0.1% Triton X-100, and 10 mM 2-mercaptoethanol (BME)) and sonicated at 90 A for 2.5 minutes of 1 second on / 5 seconds off cycles on ice. The lysate was cleared by centrifugation at 15000 rpm for 45 min at 4°C. The supernatant was loaded over 3 ml of Ni-NTA beads (Qiagen) equilibrated in lysis buffer, then washed through with wash buffer containing 50 mM Tris-HCl pH 8.0, 500 mM NaCl, 30 mM imidazole, and 10 mM BME. The recombinant SCP4 was finally eluted with elution buffer containing 50 mM Tris-HCl pH 8.0, 500 mM NaCl, 300 mM imidazole, and 10 mM BME. Protein fractions were pooled and dialyzed overnight at 4°C in a 10.0 kDa dialysis membrane (Thermo) against dialysis buffer (50 mM Tris HCl pH 7.5, 100 mM NaCl, and 10 mM BME) supplemented with 0.5 mg of TEV protease to remove the 6xHis and SUMO-tag. The protein was then loaded onto a DEAE anion exchange column (GE) equilibrated with Buffer A (50 mM Tris-HCl pH 7.5, 10 M NaCl, 10 mM BME) and eluted with a 0-100% gradient of Buffer A to Buffer B (50 mM Tris-HCl pH 7.5, 500 mM NaCl, and 10 mM BME). Peak fractions were analyzed by SDS-PAGE before the fractions containing SCP4 were pooled and dialyzed overnight in gel filtration buffer containing 25 mM Tris HCl pH 8.0, 200 mM NaCl, and 10 mM BME. The protein was finally polished using gel filtration chromatography with a Superdex 75 size exclusion column. Fractions containing SCP4 were pooled, concentrated, and flash-frozen at -80°C.

### Phosphatase activity assay

Initial rates of phosphatase activity were determined by incubating SCP4 (1µM) with various concentrations of phosphopeptide (0–100 µM) in SCP4 assay buffer (50 mM Tris-acetate pH 5.0, 10 mM MgCl_2_) in a 40µl reaction volume for 10 minutes at 37°C. 20µl of the reaction was then quenched with 40µl of BIOMOL Green reagent (Enzo Life Sciences) within a clear, flat-bottom 96-well plate. The mixtures were allowed to sit at room temperature for 30 minutes for color development before reading the absorbance signal at A_620_ in a Tecan Plate reader 200. The A_620_ values were corrected by subtracting the A_620_ values of substrate only reactions at each substrate concentration. The readings obtained were converted to the amount of phosphate released through a Phosphate standard curve determined using the Biomol green assay following the manufacturer instructions. The reaction rate was plotted for kinetic fitting to derive *k_cat_*/*K_m_*. Biological triplicates were prepared and analyzed independently to guarantee replication of experiments.

### Metabolomics analysis by liquid chromatography coupled to mass spectrometry (LC-MS)

Cells (0.6 mln cells/well) were seeded in 6 well plates after lentiviral infection with sgRNAs. On Day 5 post-infection, cells were quickly washed in PBS before adding 1 mL of ice-cold extraction solution (50% methanol, 30% acetonitrile, 20% H2O) per million cells. The cell suspension was snap frozen in liquid nitrogen. Samples were agitated using a tube rotator at 4°C for 15 min utes followed by incubation at -80°C overnight. Samples were then centrifuged at 15,000 rpm, 4°C for 10 minutes. The supernatants were collected and stored in autosampler vials at -80°C until analysis.

Intracellular extracts from five independent cell cultures were analyzed for each condition. Samples were randomized in order to avoid bias due to machine drift and processed blindly. LC-MS analysis was performed using a Vanquish Horizon UHPLC system couple to a Q Exactive HF mass spectrometer (both Thermo Fisher Scientific). Sample extracts (8 µL) were injected onto a Sequant ZIC-pHILC column (150 mm × 2.1 mm, 5 µm) and guard column (20 mm × 2.1 mm, 5 µm) from Merck Millipore kept at 45°C. The mobile phase was composed of 20 mM ammonium carbonate with 0.1% ammonium hydroxide in water (solvent A), and acetonitrile (solvent B). The flow rate was set at 200 μl/min with the previously described gradient (Mackay et al., 2015). The mass spectrometer was operated in full MS and polarity switching mode. The acquired spectra were analyzed using XCalibur Qual Browser and XCalibur Quan Browser software (Thermo Fisher Scientific) by referencing to an internal library of compounds.

## QUANTIFICATION AND STATISTICAL ANALYSIS

All the statistics tests were performed using GraphPad PRISM5 software. Error bars represent the mean plus or minus standard error of the mean, unless otherwise stated.

## DATA AND SOFTWARE AVAILABILITY

The RNA-seq data from this study have been uploaded to GEO database with accession number GSE.

---

**Figure S1.**
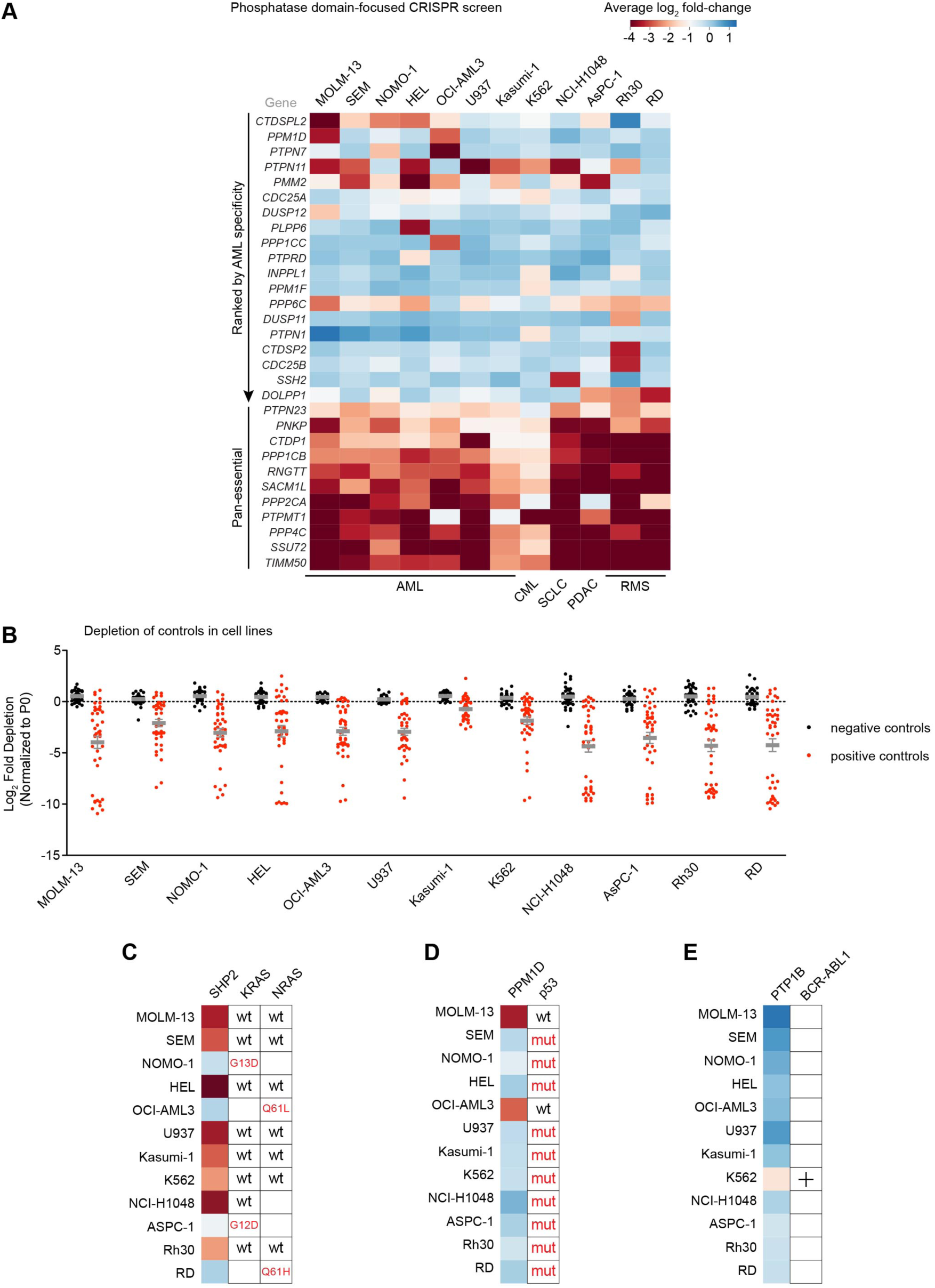
Phosphatase domain-focused CRISPR screening identifies context-specific dependencies in human cancer cell lines. **(A)** Extracted essentiality scores for phosphatases demonstrating AML-bias or pan-essentiality. Plotted is the log_2_ fold-change of sgRNA abundance during ∼11 population doublings. The effects of individual sgRNAs targeting each domain were averaged. AML, acute myeloid leukemia; CML, chronic myeloid leukemia; SCLC, small cell lung cancer; PDAC, pancreatic ductal adenocarcinoma; RMS, rhabdomyosarcoma. **(B)** The log_2_ fold-changes of individual sgRNAs serving as negative (black) or positive (red) controls in the screen. **(C–E)** Extracted essentiality scores for selected phosphatases. Plotted is the log_2_ fold-change of sgRNA abundance during ∼11 population doublings. The effects of individual sgRNAs targeting each domain were averaged. Also shown is the relevant oncogene/tumor suppressor status for each cell line.

**Figure S2.**
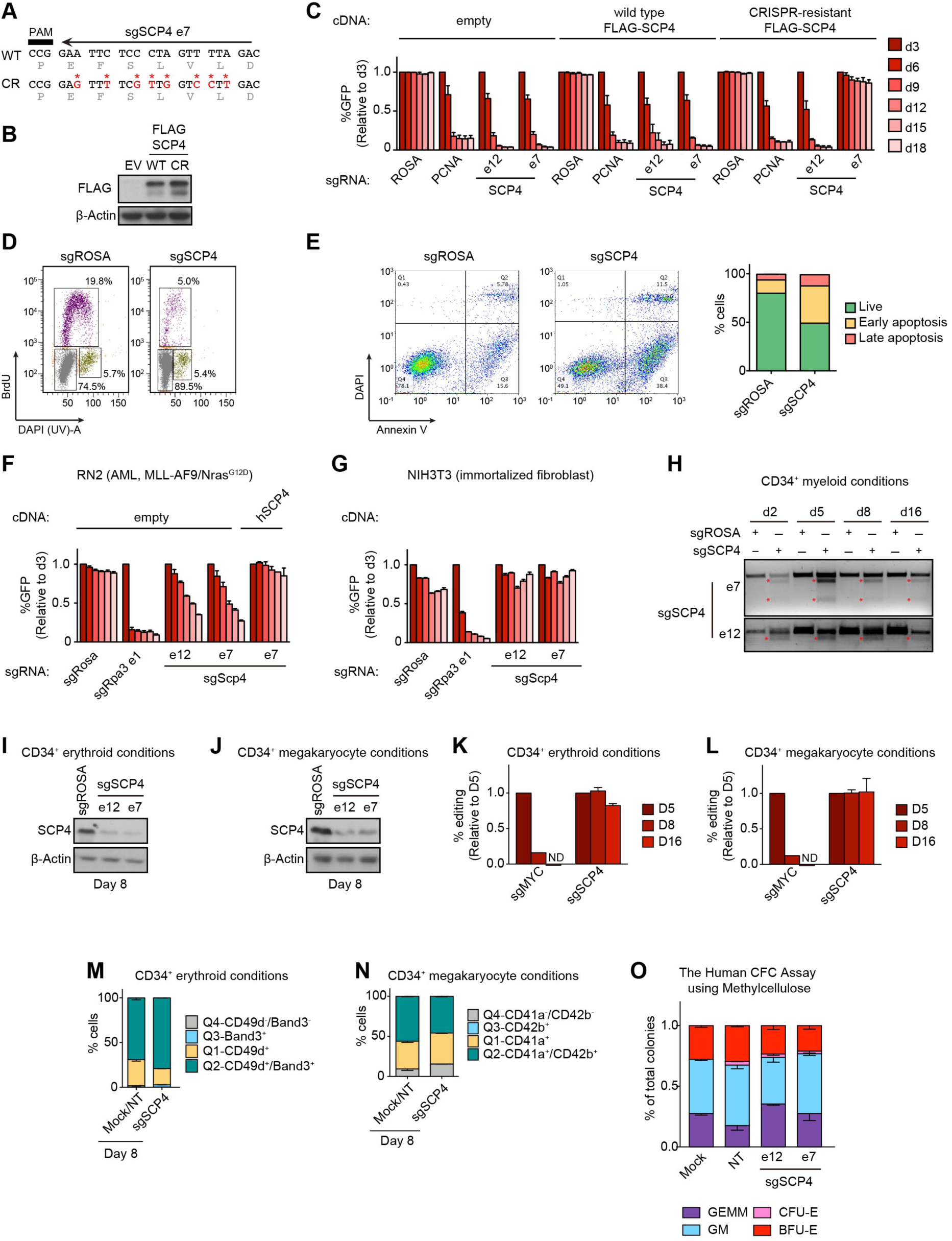
SCP4 is an acquired dependency in human AML. **(A)** Design of CRISPR-resistant mutant of SCP4. **(B)** Western blot of whole-cell lysates from MOLM-13 cells stably expressing empty vector (EV), wild type (WT), or CRISPR-resistant (CR) FLAG-SCP4. **(C)** Competition-based proliferation assay in MOLM-13 cells stably expressing empty vector, wild type, or CRISPR-resistant FLAG-SCP4 infected with the indicated sgRNAs. n = 3. **(D)** Representative flow cytometry analysis of BrdU incorporation and DNA content to infer cell cycle stage in MOLM-13 cells on day 5 post-infection with the indicated sgRNAs. **(E)** (Left) Representative flow cytometry analysis of DAPI (indicating permeable dead cells) and annexin-V staining (a pre-apoptotic cell marker) in MOLM-13 cells on day 5 post-infection with the indicated sgRNAs. (Right) Quantification of live and apoptotic cells. n = 3. **(F–G)** Competition-based proliferation assays in the murine cell lines infected with the indicated sgRNAs. Rescue in the RN2 cells stably expressing human SCP4 is shown. n=3. **(H)** Surveyor Assay analysis of indels presence during in CD34^+^ cells electroporated with Cas9 loaded with the indicated sgRNAs over the course of culturing in myeloid conditions. **(I–J)** Western blot of whole-cell lysates from CD34^+^ cells electroporated with Cas9 loaded with the indicated sgRNAs, day 8 post-electroporation in erythroid (I) and megakaryocyte (J) conditions. **(K–L)** RT-qPCR analysis of indels presence in CD34^+^ cells electroporated with Cas9 loaded with the indicated sgRNAs over the course of culturing in erythroid (K) and megakaryocyte (L) conditions. The effects of individual sgRNAs for SCP4 were averaged. n = 4. **(M–N)** Quantification of the flow cytometry analysis of erythroid (M) and megakaryocyte (N) differentiation of CD34^+^ cells electroporated with Cas9 loaded with the indicated sgRNAs, day 8 post-electroporation, culturing in erythroid (M) and megakaryocyte (N) conditions. The effects of individual negative controls and sgRNAs for SCP4 were averaged. n = 4. **(O)** The Human Colony Forming Cell (CFC) assay using methylcellulose. GEMM, colony forming unit-granulocyte, erythrocyte, macrophage, megakaryocyte; GM, colony forming unit-granulocyte, macrophage; CFU-E, colony forming unit-erythroid; BFU-E, burst forming unit-erythroid. All bar graphs represent the mean ± SEM. All sgRNA experiments were performed in Cas9- expressing cell lines. ‘e’ refers to the exon number targeted by each sgRNA. The indicated sgRNAs were linked to GFP. GFP^+^ population depletion indicates loss of cell fitness caused by Cas9/sgRNA-mediated genetic mutations. ROSA, Mock, and NT, negative controls; PCNA, Rpa3, and MYC, positive controls.

**Figure S3.**
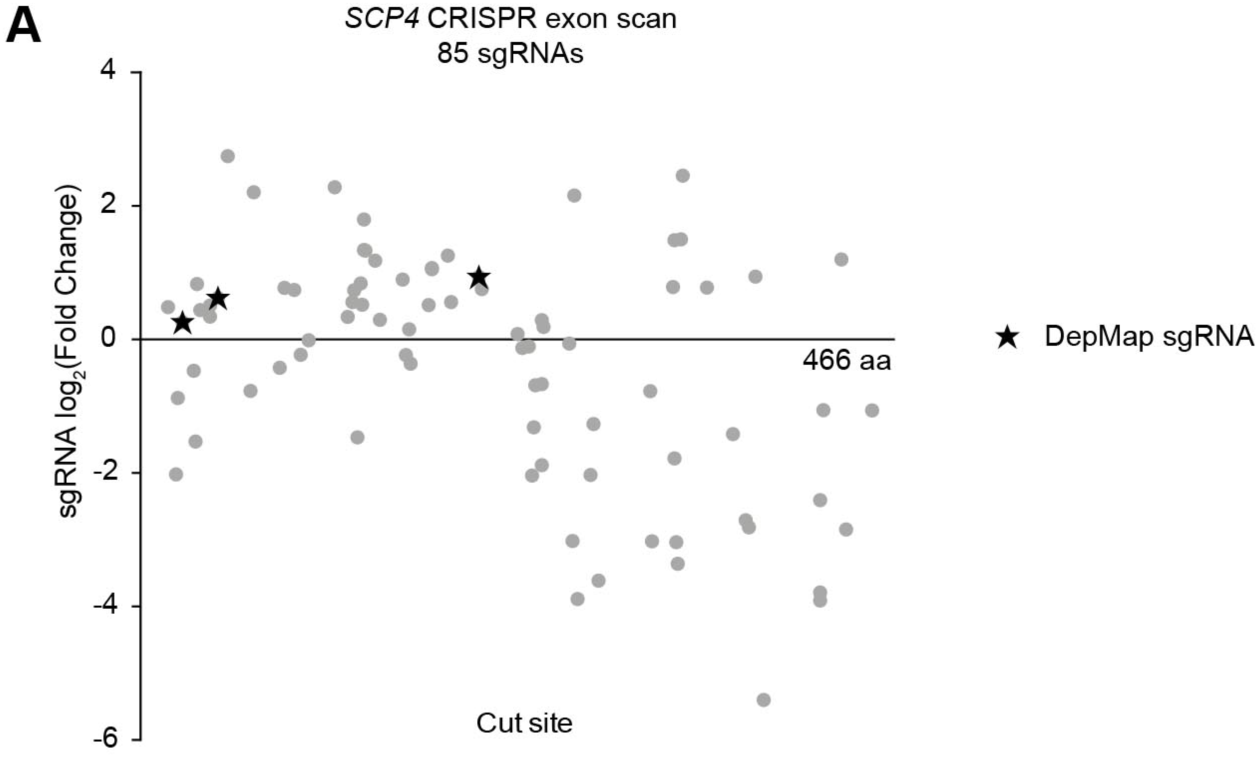
The catalytic phosphatase function of SCP4 is essential in AML. **(A)** The CRISPR- scan of SCP4 with all the possible sgRNAs. Deep sequencing-based measurement of the impact of 85 SCP4 sgRNAs on the proliferation of MOLM-13 cells.

**Figure S4.**
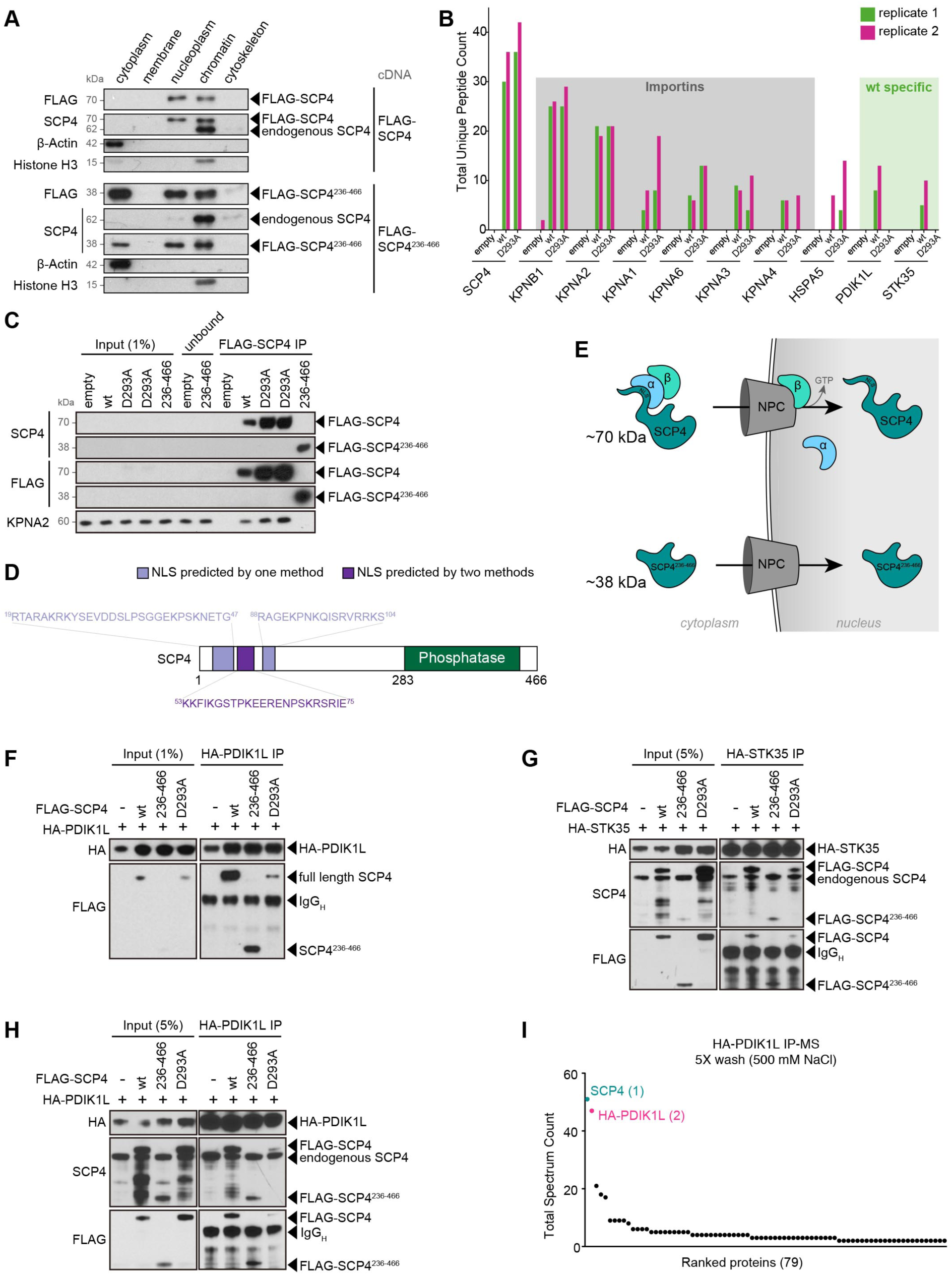
SCP4 directly interacts with two kinase homologs STK35 and PDIK1L. **(A)** Western blot of the indicated fractions from MOLM-13 cells stably expressing FLAG-SCP4^WT^ and FLAG-SCP4^236-466^. **(B)** Total unique peptide counts for the top hits detected by MS in two independent biological replicates. **(C)** Immunoprecipitation followed by Western blotting performed with the indicated antibodies. The nuclear lysates were prepared from the human MOLM-13 cells stably expressing empty vector, FLAG-SCP4^WT^, FLAG-SCP4^236-466^, catalytic mutant FLAG-SCP4^D293A^. The flow-through was analyzed to ensure efficient binding of the FLAG-tagged constructs (loaded as “unbound”). **(D)** Nuclear localization signals (NLS) predictions for SCP4 amino acid sequence performed by two different methods (Nguyen Ba et al., 2009; Kosugi et al., 2009). **(E)** Model of full length and truncated SCP4 transport into the nucleus. **(F–H)** Immunoprecipitation followed by Western blotting performed with the indicated antibodies. The whole-cell lysate was prepared from HEK293T 24 hours post-transfection with the indicated constructs (F). The nuclear lysates were prepared from the human AML cell line MOLM-13 stably expressing the indicated constructs (G, H). ‘-’, empty vector; WT, wild type FLAG-SCP4; 236-466, FLAG-SCP4^236-466^; D293A, catalytic mutant FLAG-SCP4^D293A^, IP, immunoprecipitation. Note: degradation bands appear in the WT and D293A input at ∼50 kDa and at ∼40 kDa and the WT IP at ∼40 kDa. **(I)** Total spectrum counts (TSC) for HA-PDIK1L and endogenous SCP4 detected by mass spectrometry on Pierce Anti-HA Magnetic Beads after stringent washes with washing buffers containing 500 mM NaCl.

**Figure S6.**
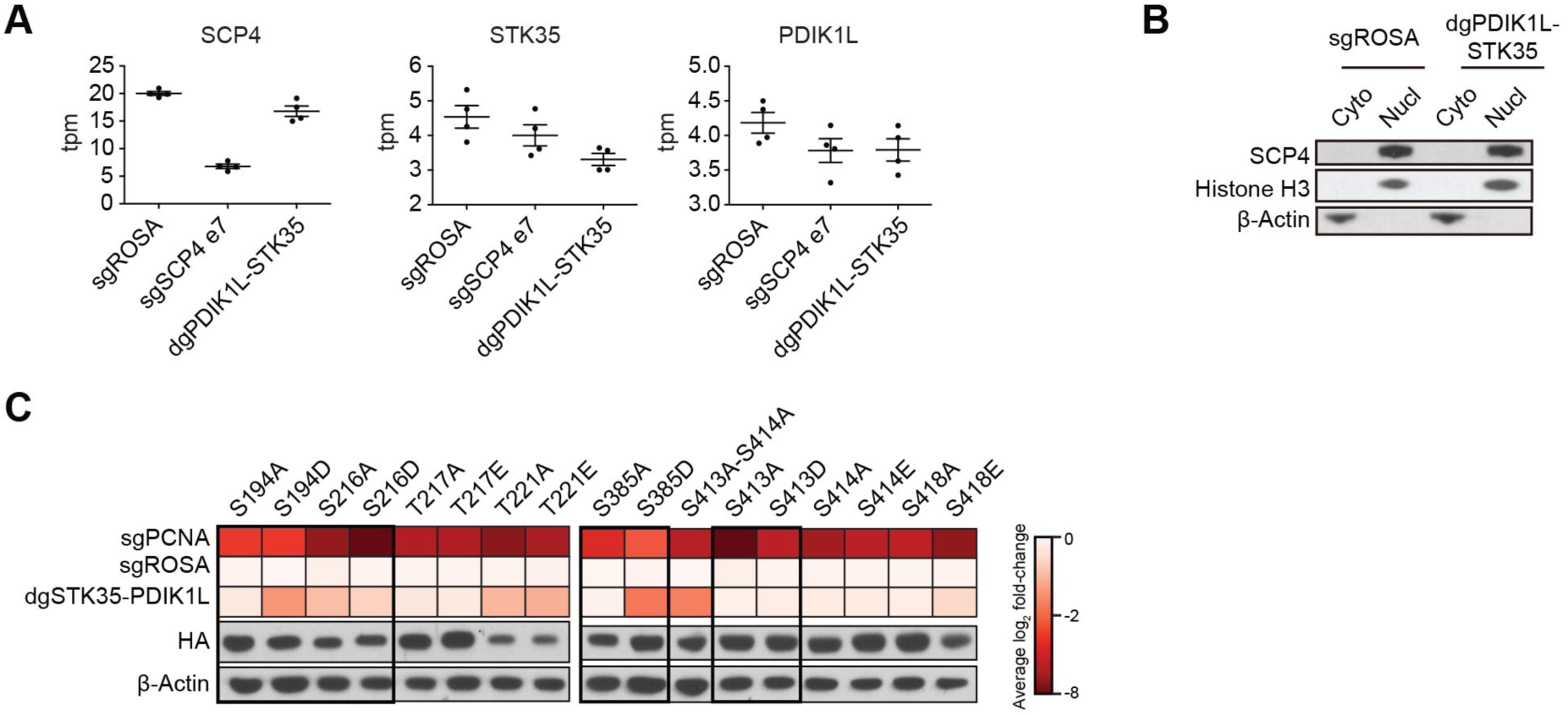
SCP4 functions upstream of STK35/PDIK1L to support AML proliferation. **(A)** RNA-Seq transcripts per million (tpm) data for SCP4, STK35, and PDIK1L from MOLM-13 cells on day 5 post-infection with the indicated sgRNAs. Plotted are data from each individual RNA-Seq replicate, the mean ± SEM. **(B)** Western blot of cytoplasm (Cyto) and nucleus (Nucl) fractions of MOLM-13 cells on day 5 post-infection with the indicated sgRNAs. STK35/PDIK1L simultaneous depletion was assessed by the loss of fitness during 18 days in culture showed in Figure 5A. **(C)** Summary of competition-based proliferation assays in MOLM-13 cells stably expressing CRISPR-resistant HA-PDIK1L or HA-STK35 constructs harboring the indicated amino acid substitutions infected with the indicated sgRNAs. Plotted is the fold change (log2) of sgRNA^+^/GFP^+^ cells after 18 days in culture (average of triplicates). Below are shown Western blots of whole-cell lysates of the cells stably expressing the indicated constructs. All sgRNA experiments were performed in Cas9-expressing cell lines. ‘e’ refers to the exon number targeted by each sgRNA. ‘dg’ refers to the bi-cistronic vector for simultaneous targeting of STK35 and PDIK1L. Negative fold-change indicates loss of cell fitness caused by Cas9/sgRNA-mediated genetic mutations.

**Figure S7.**
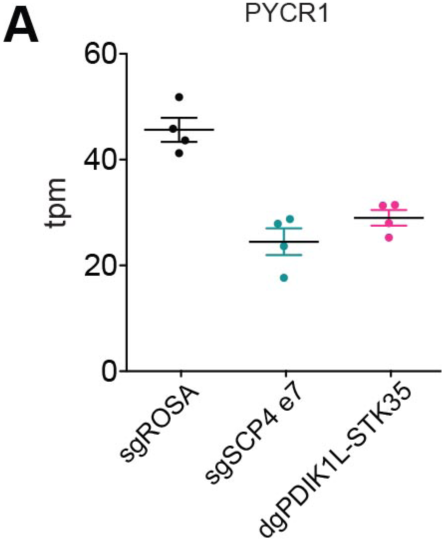
SCP4-kinase complex supports metabolic homeostasis of AML cells. **(A)** RNA-Seq transcripts per million (tpm) data for *PYCR1* from MOLM-13 cells on day 5 post-infection with the indicated sgRNAs. Plotted are data from each individual RNA-Seq replicate, the mean ± SEM. All sgRNA experiments were performed in Cas9-expressing cell lines. ‘e’ refers to the exon number targeted by each sgRNA. ‘dg’ refers to the bi-cistronic vector for simultaneous targeting of STK35 and PDIK1L.

